# Global characterization of copy number variants in epilepsy patients from whole genome sequencing

**DOI:** 10.1101/199224

**Authors:** Jean Monlong, Simon L. Girard, Caroline Meloche, Maxime Cadieux-Dion, Danielle M. Andrade, Ron G. Lafreniere, Micheline Gravel, Dan Spiegelman, Alexandre Dionne-Laporte, Cyrus Boelman, Fadi F. Hamdan, Jacques L. Michaud, Guy Rouleau, Berge A. Minassian, Guillaume Bourque, Patrick Cossette

## Abstract

Epilepsy will affect nearly 3% of people at some point during their lifetime. Previous copy number variants (CNVs) studies of epilepsy have used array-based technology and were restricted to the detection of large or exonic events. In contrast, whole-genome sequencing (WGS) has the potential to more comprehensively profile CNVs but existing analytic methods suffer from limited accuracy. We show that this is in part due to the non-uniformity of read coverage, even after intra-sample normalization. To improve on this, we developed PopSV, an algorithm that uses multiple samples to control for technical variation and enables the robust detection of CNVs. Using WGS and PopSV, we performed a comprehensive characterization of CNVs in 198 individuals affected with epilepsy and 301 controls. For both large and small variants, we found an enrichment of rare exonic events in epilepsy patients, especially in genes with predicted loss-of-function intolerance. Notably, this genome-wide survey also revealed an enrichment of rare non-coding CNVs near previously known epilepsy genes. This enrichment was strongest for non-coding CNVs located within 100 Kbp of an epilepsy gene and in regions associated with changes in the gene expression, such as expression QTLs or DNase I hypersensitive sites. Finally, we report on 21 potentially damaging events that could be associated with known or new candidate epilepsy genes. Our results suggest that comprehensive sequence-based profiling of CNVs could help explain a larger fraction of epilepsy cases.

**Author summary:** Epilepsy is a common neurological disorder affecting around 3% of the population. In some cases, epilepsy is caused by brain trauma or other brain anomalies but there are often no clear causes. Genetic factors have been associated with epilepsy in the past such as rare genetic variations found by linkage studies as well as common genetic variations found by genome-wide association studies and large copy-number variants. We sequenced the genome of *∼*200 epilepsy patients and *∼*300 healthy controls and compared the distribution of deletion (loss of a copy) and duplication (additional copy) of genomic regions. Thanks to the sequencing technology and a new method that takes advantage of the large sample size, we could compare the distribution of small copy-number variants between epilepsy patients and controls. Overall, we found that small variants are also associated with epilepsy. Indeed, the genome of epilepsy patients had more exonic copy-number variants, especially when rare or affecting genes with predicted loss-of-function intolerance. Focusing on regions around genes that have been previously associated with epilepsy, we also found more non-coding variants in epilepsy patients, especially deletions or variants in regulatory regions. Finally, we provide a list of 21 regions in which we found likely pathogenic variants.

## 1 Introduction

Structural variants (SVs) are defined as genetic mutations affecting more than 50 base pairs and encompass several types of rearrangements: deletion, duplication, novel insertion, inversion and translocation. Deletions and duplications, which affect DNA copy number, are collectively known as copy number variants (CNVs). SVs arise from a broad range of mechanisms and show a heterogeneous distribution of location and size across the genome [1–3]. Numerous diseases are caused by SVs with a demonstrated detrimental effect [4, 5]. While cytogenetic approaches and array-based technologies have been used to identify large SVs, whole-genome sequencing (WGS) has the potential to uncover the full range of SVs both in terms of type and size [6,7]. SV detection methods that use read-pair and split read information [8] can detect deletions and duplications but most CNV-focused approaches look for an increased or decreased read coverage, the expected consequence of a duplication or a deletion. Coverage-based methods exist to analyze single samples [9], pairs of samples [10] or multiple samples [11–13] but the presence of technical bias in WGS remains an important challenge. Indeed, various features of sequencing experiments, such as mappability [14, 15], GC content [16], replication timing [17], DNA quality and library preparation [18], have a negative impact on the uniformity of the read coverage [19].

Epilepsy is a common neurological disorder characterized by recurrent and unprovoked seizures. It is estimated that up to 3% of the population will suffer from a form of epilepsy at some point during their lifetime. Although the disease presents a strong genetic component that can be as high as 95%, typical “monogenic” epilepsy is rare, accounting for only a fraction of cases [20,21]. Genetic factors have been associated with epilepsy in the past such as rare genetic variations found by linkage studies as well as common genetic variations found by genome-wide association studies [22, 23] For example, a meta-analysis combining multiple epilepsy cohorts found positive associations with the disease [24], the strongest in *SCN1A*, a gene already associated with the genetic mechanism of the disease via linkage studies and subsequent sequencing [25] or more recently as harboring de novo variants [26]. Thanks to array-based technologies, surveys of large CNVs (*>*50 Kbp) first associated CNVs in genomic hotspots such as 15q11.2 and 16p13.11 with generalized epilepsy [27, 28]. Other studies have further shown the importance of large and *de novo* CNVs as well as identified a few associations with specific genes [29–34]. Rare genic CNVs were typically found in around 10% of epilepsy patients [30, 34, 35] and CNVs larger than 1 Mbp were significantly enriched in patients compared to controls [33, 35–37]. Unfortunately, small CNVs and other types of SVs could not be efficiently or consistently detected using these technologies, hence much remains to be done.

To more comprehensively characterize the role of CNVs in epilepsy, we performed whole-genome sequencing of epileptic patients from the Canadian Epilepsy Network (CENet), the largest WGS study on epilepsy to date. In the present study, we assessed the frequency of CNVs in epileptic individuals using 198 unrelated patients and 301 healthy individuals. Using this data, we showed that technical variation in WGS remains problematic for CNV detection despite state-of-the-art intra-sample normalization. To correct for this and to maximize the potential of the CENet cohorts, we developed a population-based CNV detection algorithm called PopSV. Our method uses information across samples to avoid systematic biases and to more precisely detect regions with abnormal coverage. Using two public WGS datasets [38, 39], and additional orthogonal validation, we showed that PopSV outperforms other analytical methods both in terms of specificity and sensitivity, especially for small CNVs. Using this tool, we built a comprehensive catalog of CNVs in the CENet epilepsy patients and studied the properties of these potentially damaging structural events across the genome.

## 2 Results

### 2.1 Technical bias in read coverage

We sequenced the genomes of 198 unrelated individuals affected with epilepsy and 301 unrelated healthy controls. Because CNV detection relies on read coverage we first investigated the presence of technical bias and the value of standard corrections and filters (e.g. GC correction, mappability filtering). The genome was fragmented in 5 Kb bins and we counted the number of uniquely mapped reads in each bin. In contrast to simulated datasets, we found that the inter-sample mean coverage in each bin varied between genomic regions even after stringent corrections and filters (Fig 1a). Supporting this observation, the bin coverage variance across samples was also lower than expected and varied between regions (Fig S1a). We also observed experiment-specific biases. In particular, some samples consistently had the highest, or the lowest, coverage across large portions of the genome (Fig S1b). These observations were not unique to our data and could also be observed in two public WGS datasets, and persisted even after correcting the GC bias and mappability using the more elaborate model from the QDNAseq pipeline [40] (Fig S2). Our results across multiple samples suggest that existing GC bias and mappability corrections [40] cannot correct completely the technical variation in read coverage. This fluctuation of coverage has implications for CNV detection approaches that assume a uniform distribution [9, 10, 41] after standard bias correction and will lead to false positives.

**Fig 1:**
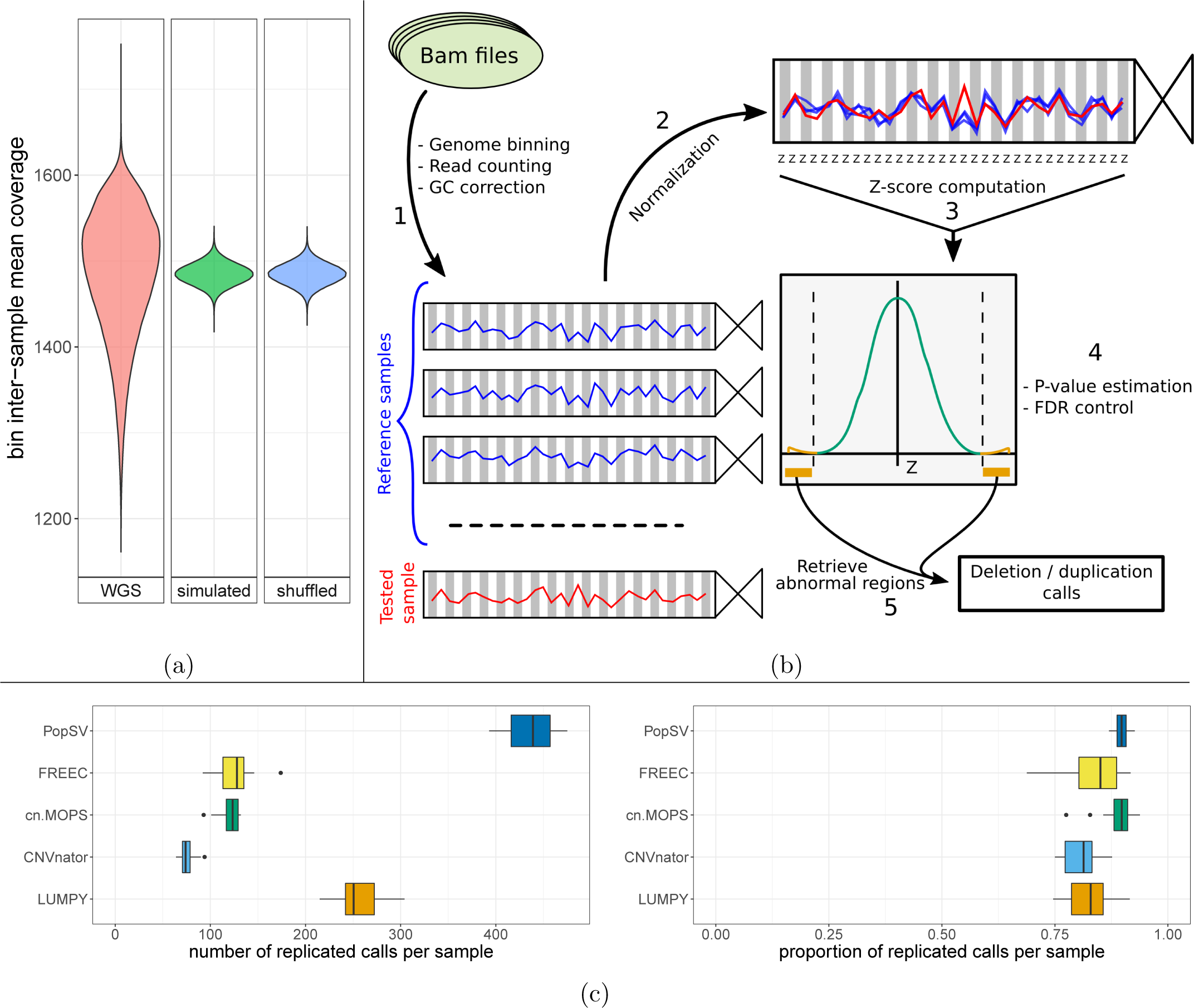
PopSV approach. a) Technical bias across the genome remains after stringent correction and filtering. The distribution of the bin inter-sample mean coverage in the epilepsy cohort (red) is compared to null distributions (blue: bins shuffled, green: simulated normal distribution). b) PopSV approach. First the genome is fragmented and reads mapping in each bin are counted for each sample and GC corrected (1). Next, coverage of the sample is normalized (2) and each bin is tested by computing a Z-score (3), estimating p-values (4) and identifying abnormal regions (5).c) Number and proportion of calls from a twin that was replicated in the other monozygotic twin.

### 2.2 CNV detection with PopSV

To better control for technical bias, we developed PopSV, a new SV detection method. PopSV uses read depth across the samples to normalize coverage and detect change in DNA copy number (Fig 1b). The normalization step here is critical since most approaches will fail to give accept- able normalized coverage scores (Fig S1b). Moreover, with global median/variance adjustment or quantile normalization, the remaining subtle experimental variation impairs the abnormal coverage test (Fig S3a). The targeted normalization used by PopSV was found to have better statistical properties (Fig S3b). In order to assess the performance of our tool, we compared it to several algorithms [8–11] using a dataset that included monozygotic twins and also performed experimental validation of different types of predicted CNVs in the epilepsy cohort (see below). We found that PopSV performed as well or better in different aspects. First, for several algorithms, a large proportion of the detected events in a typical sample were also identified in almost all samples (60% of the calls found in *>*95% of the samples, Fig S4). PopSV’s calls were better distributed across the frequency spectrum, hence more informative as we expect the relative frequency of disease-related variants to be rare. In addition, the pedigree structure was more accurately recovered when the CNVs were used to cluster the individuals in the Twins dataset (Fig S5). The agreement with the pedigree was computed by the Rand index after clustering the individuals with three hierarchical clustering approaches (see Supplementary Information). Looking at the replication between 10 pairs of monozygotic twins, PopSV detected more replicated CNVs compared to other methods, while maintaining similar replication rates (Fig 1c). The CNV calls were further filtered with gradually more stringent significance thresholds and PopSV remained superior in term of number of replicated calls (Fig S6). When investigating the overlap of calls between different methods, we noticed that PopSV was better recovering calls from CNVnator [9], FREEC [10], cn.MOPS [11] or LUMPY [8], especially if found by two or more methods (Fig S7). For example, around 92% of the CNVs called by other methods were also found by PopSV when focusing on calls found in at least two methods. Similar results were also obtained in a cancer dataset where we looked for replicated germline CNVs in the paired tumor (Fig S8). Finally, we repeated the twin analysis using 500 bp bins and observed high consistency with the 5 Kbp calls (Fig S9). These results suggest that PopSV can accurately detect around 75% of events that are as large as half the bin size used (see Supplementary Information).

### 2.3 CNVs in the CENet cohorts and experimental validation

Having demonstrated the quality of the PopSV calls, we applied our tool to the epilepsy and control cohorts. The epilepsy cohort comprises 198 individuals diagnosed with either generalized (n=160), focal (n=32) or unclassified (n=6) epilepsy. CNVs ranged from 5 Kbp to 3.2 Mbp with an average size of 9.98 Kbp. We observed an average of 870 CNVs per individual accounting for 8.7 Mb of variant calls (Fig 2a). This is around 9 times more variants and considerably smaller than in typical array-based studies [42, 43], such as the previous epilepsy surveys [30, 31, 34, 35], although a similar size distribution was previously obtained using denser arrays [4] but were never applied to epilepsy (Fig S10). Next, we annotated each variant using four public SV databases [13, 44–46] as well as an internal database of the germline calls from PopSV in the two public datasets used earlier (see Supplementary Information). For each CNV, we derived the maximum frequency across these databases and defined as rare any region consistently annotated in less than 1% of the individuals (Fig 2b). In total, we identified 12,480 regions with rare CNVs in the epilepsy cohort including: 8,022 (64.3%) with heterozygous deletions, 21 (0.2%) with homozygous deletions and 4,850 (38.9%) with duplications. Although the overall amount of rare CNVs was not higher in epilepsy patients, the proportion of deletion was significantly higher compared to controls (*χ*^2^ test: P-value 10^−7^). Next, we selected 151 CNVs and further validated them using a Taqman CNV assay and Real-Time PCR. To explore PopSV’s performance across different CNV profiles, we selected variants of different types, sizes and frequencies. We found that the calls were concordant in 90.7% of the cases (Table 1 and S2). As expected, the estimated false positive rate was slightly higher for rare or smaller variants (12.1% for rare CNVs; 15.1% for CNV *<*20 Kbp). Furthermore, we noted that calls supported by both PopSV and LUMPY (when available) had a similar validation rate as calls found by PopSV only (86.2% and 87.5% respectively).

**Table 1:** Real-Time PCR validation rates of PopSV calls. Number and proportion of regions validated for CNVs of different types, sizes and frequencies.

|  | Region | Validation rate |
| --- | --- | --- |
| Total | 151 | 0.907 |
| CNV type |  |  |
| Deletion | 102 | 0.902 |
| Duplication | 49 | 0.918 |
| Frequency in databases |  |  |
| 0 | 26 | 0.923 |
| (0, 0.01] | 24 | 0.833 |
| (0.01, 1] | 101 | 0.921 |
| Carrier in CENet cohorts |  |  |
| 1 | 21 | 0.857 |
| 2 | 19 | 0.947 |
| > 2 | 111 | 0.910 |
| Size (Kbp) |  |  |
| < 20 | 73 | 0.849 |
| (20, 100] | 38 | 0.974 |
| > 100 | 40 | 0.950 |

**Fig 2:**
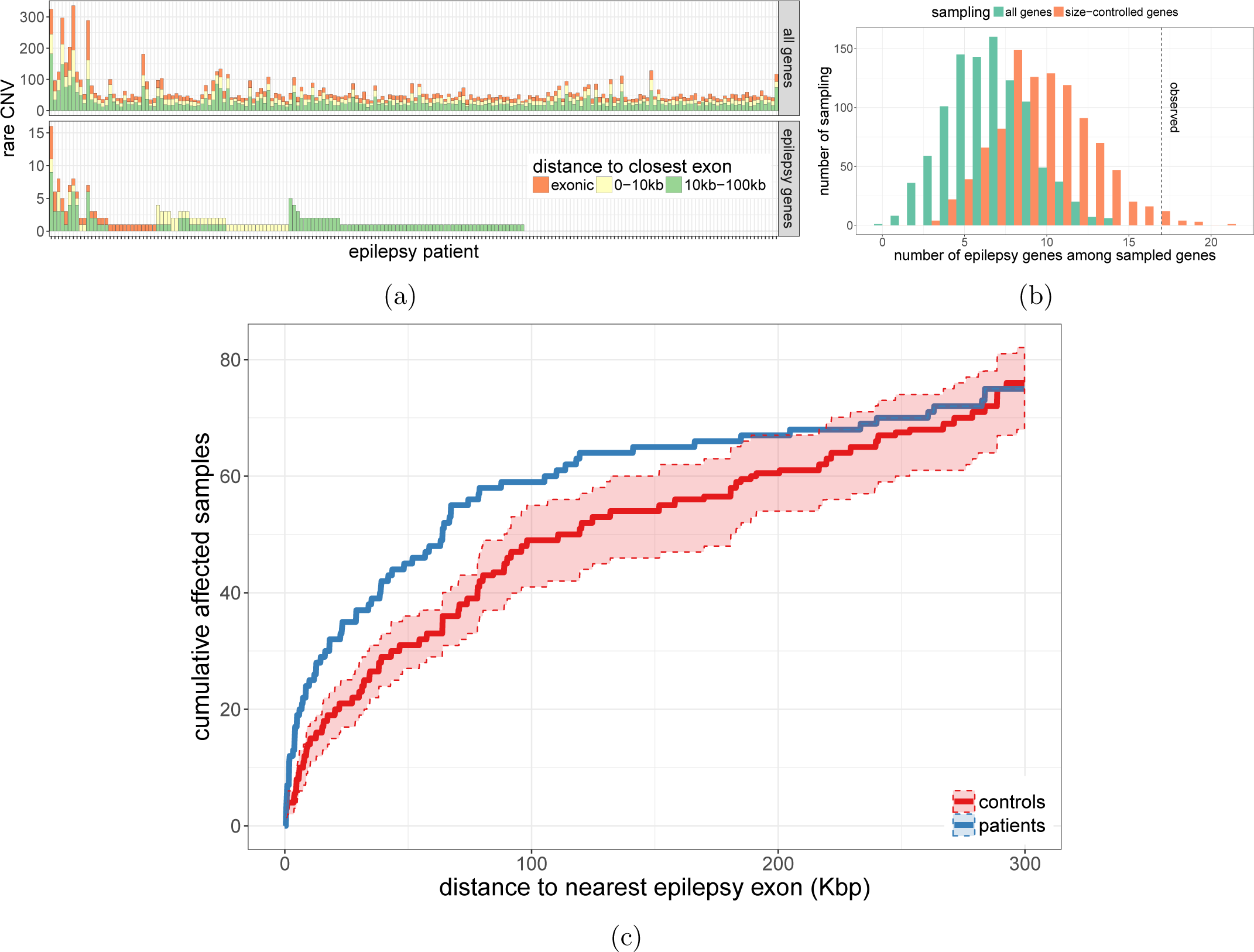
CNVs in the epilepsy and control cohorts. a) Regions with a CNV in each epilepsy patient. b) Each CNV in the CNV catalog of the epilepsy and control cohorts was annotated with its maximum frequency in five CNV databases. c) Enrichment in exonic sequence for all CNVs (left) and rare CNVs (right), larger than 50 Kbp (top) or smaller than 50 Kbp (bottom). The fold-enrichment (y-axis) represents how many CNVs overlap coding sequences compared to control regions randomly distributed in the genome.

### 2.4 CNV enrichment in exonic regions

To assess the role of CNVs in the pathogenic mechanism of epilepsy, we evaluated the prevalence of exonic CNVs in our epileptic cohort compared with healthy controls. First, focusing on CNVs larger than 50 Kbp, we found no difference between epileptic patients and controls (Fig 2c). As expected, we observed fewer CNVs overlapping exonic sequence than expected by chance but similar levels for both groups. The number of CNVs overlapping exonic sequences of genes intolerant to loss-of-function mutations [47] was even lower. Interestingly, the coding regions of those genes were significantly more affected by CNVs in epileptic patients compared with controls (permutation P-value*<*0.001, Figs 2c and S11). Because they are more likely pathogenic and of greater interest, we performed the same analysis using rare CNVs only. Here, we observed the increased exonic burden described previously for large rare CNVs [35–37]. In contrast to previous studies, we could also detect and compare small CNVs (*<*50 Kbp) in epileptic patients and healthy controls. We found similar enrichment patterns than for large CNVs (Fig 2c and S11), suggesting that small rare exonic CNVs are also associated with epilepsy. Indeed, there was no significant difference between epileptic patients and controls when considering all small CNVs and all genes. The exonic enrichment was significant for genes with predicted loss-of-function intolerance and for rare variants (permutation P-value*<*0.001, Fig 2c and S11). In both cohorts, most of the rare exonic CNVs were private, i.e. present in only one individual. However, we observed that rare exonic CNVs were less likely private in the epileptic patients (permutation P-value*<*0.001, Fig S12a). We replicated this result using only individuals with a similar population background (French-Canadians, Fig S12b). Overall we concluded that rare CNVs were not only enriched in exons but also affected exons more recurrently in the epilepsy cohort as compared to controls.

### 2.5 CNV enrichment in and near epilepsy genes

We then sought to evaluate if there was an excess of CNVs disrupting epilepsy-related genes or nearby functional regions. We first retrieved genes whose exons were hit by rare deletions or duplications and evaluated how many were known epilepsy genes based on a list of 154 genes previously associated with epilepsy [48] (Fig 3a). Because epilepsy genes tend to be large, we controlled for the gene size when testing for enrichment (Fig S13a). In the epilepsy cohort only, we noted a clear enrichment for epilepsy genes hit by rare deletions (Fig S13b). Moreover, the enrichment became stronger for rare CNVs. For instance, the exons of 921 genes were disrupted in the epilepsy cohort when considering deletions completely absent from the public and internal databases, 17 of which were epilepsy genes (P-value 0.015, Fig 3b). In addition, we observed significantly more epilepsy patients with a rare non-coding CNV close to an epilepsy gene compared to control individuals (Fig S14a). Interestingly, this enrichment was stronger for non-coding deletions (Fig S14b). We further explored the distribution of rare non-coding deletions by testing each epilepsy gene for a difference in mutation load between patients and controls. The *GABRD* gene had the strongest and only nominally significant association with four non-coding deletions among the 198 epileptic patients and none in the 301 controls. *GABRD* encodes the delta subunit of the gamma-aminobutyric acid A receptor and has been associated with juvenile myoclonic epilepsy [49]. In our cohort, two of the four patients with a rare non-coding deletion close to *GABRD* had been diagnosed with this syndrome, including one patient with a 2.7 Kbp deletion located only 3 Kbp upstream of *GABRD*’stranscription start site (Fig S16a). Although none survived multiple testing correction, we noted that the strongest associations were all in the direction of a higher mutation load in the epilepsy cohort rather than in the control cohort.

**Fig 3:**
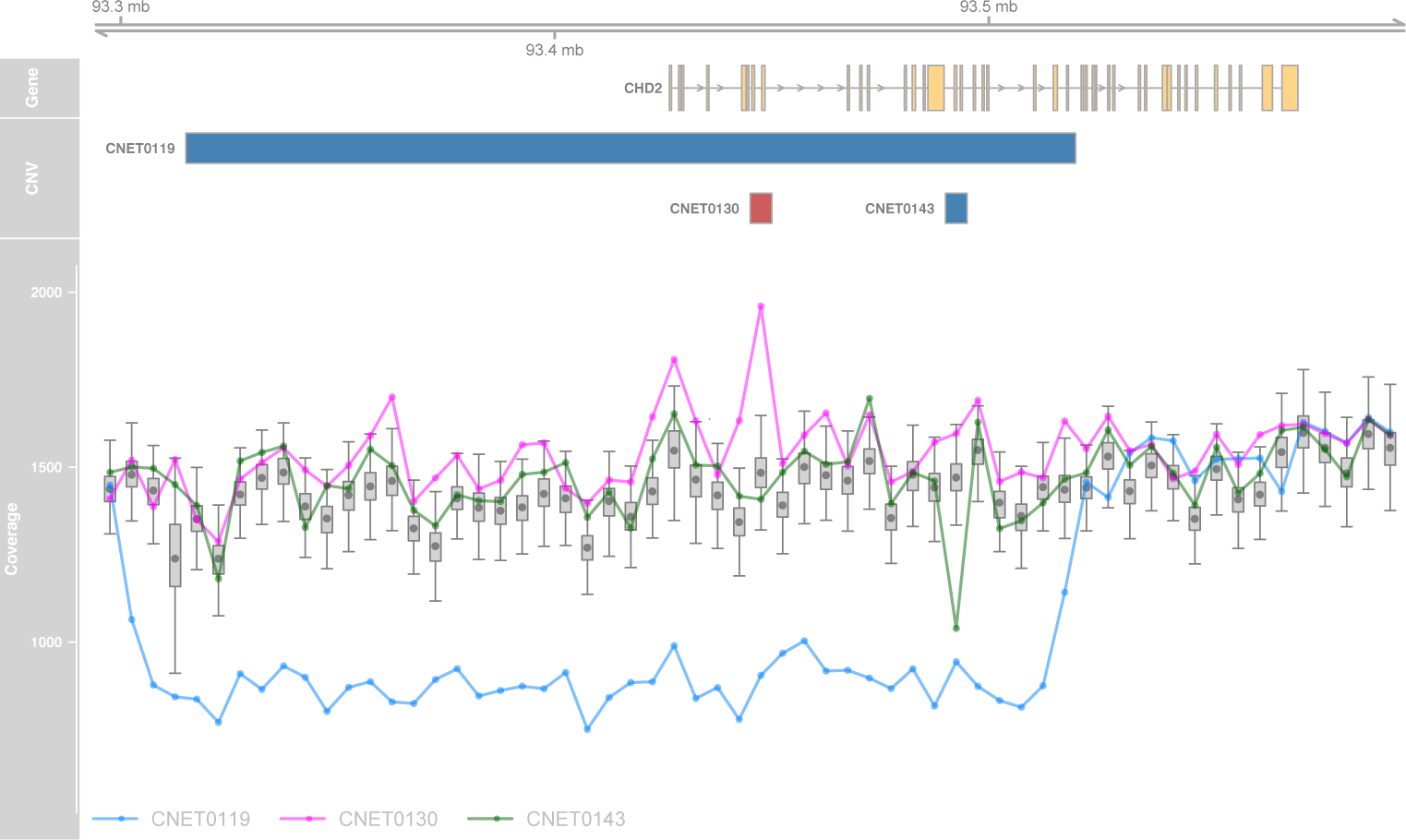
CNVs and epilepsy genes. a) Number of rare CNVs in or close to exons of protein-coding genes (top) or epilepsy genes (bottom), in the epilepsy cohort. b) Number of epilepsy genes hit by exonic deletions in the epilepsy cohort and never seen in the public and internal databases (dotted line), compared to the expected distribution in all genes and size-matched genes (histograms). c) Rare non-coding CNVs in functional regions near epilepsy genes. The graph shows the cumulative number of individuals (y-axis) with a rare non-coding CNV located at X Kbp or less (x-axis) from the exonic sequence of a known epilepsy gene. We used CNVs overlapping regions functionally associated with the epilepsy gene (eQTL or promoter-associated DNase site).

To get a better idea of the functional regions close to epilepsy genes, we retrieved their associated eQTLs in the GTEx database [50] and the DNase hypersensitivity sites associated with their promoter regions [51]. Notably, focusing on rare non-coding CNVs overlapping these functional regions, the enrichment in epileptic patients was greatly strengthened and clearly present up to 100 Kbp from an epilepsy gene (Kolmogorov-Smirnov test: P-value 9 *×* 10^*?*^5^^, Fig 3c). Comparing epilepsy patients and controls, the odds ratio of having such a CNV at a distance of 100 Kbp or less from an exon was 1.33 and gradually increased the closer to the exon (2.9 for CNVs at 5 Kbp or less, Fig S15). These non-coding CNVs were rare even in the epileptic cohort, but collectively represented an important fraction of affected patients. While 20 patients (10.1%) had exonic CNVs in epilepsy genes that were not seen in any control or in the public and internal databases, this number rose to 57 patients (28.8%) when counting non-coding CNVs in functional regions located at less than 100 Kbp of an epilepsy gene. These non-coding CNVs were never seen in the controls nor the CNV databases and overlap with annotated enhancer of epilepsy genes. Although their functional impact remains putative, we believe these CNVs to be of high-interest for the identification of disease causing genes. Among these CNVs of high-interest, a duplication of a regulatory region 5 Kbp downstream of *CSNK1E* was detected and validated in two different patients but absent from our controls and the public and internal databases (Fig S16b). Another example is a short deletion of an extremely conserved region downstream of *FAM63B*, detected in one patient and overlapping expression QTLs for this epilepsy gene (Fig S16c).

### 2.6 Putatively pathogenic CNVs

Next, we used an array of criteria to select the rare CNVs (less than 1% in 301 controls) with the highest disruptive potential in the epilepsy cohort. Priority was given to exonic CNVs in genes already known to be associated with epilepsy. For CNVs in other genes, we also prioritize recurrent variants and deletions in genes highly intolerant to loss-of-function mutations. In total, we identified 21 such putative pathogenic CNVs (Tables 2-3 and Table S3). Out of these, 8 directly affected a gene previously associated with epilepsy [48] (Table 2). In particular, we identified a deletion resulting in the loss of more than half of the *DEPDC5* gene in a patient affected with partial epilepsy. A number of point mutations have previously been reported in this gene for the same condition [52, 53]. We also identified two deletions and one duplication in *CHD2* gene (see Fig 4). The first deletion is large and affects a major portion of the gene while the second is a small 4.6 Kbp deletion of exon 13, the last exon of *CHD2*’s second isoform (Fig S17). No exon-disruptive CNVs were reported in any individuals from the control cohort. This gene was previously associated with patients suffering from photosensitive epilepsy [54]. Interestingly, all three patients carrying the CNVs in *CHD2* have been diagnosed with eyelid myoclonia epilepsy with absence, the same diagnosis that was largely enriched in the Galizia *et al.* study. Other known epilepsy genes affected by deletions include *LGI1* and the 15q13.3 region.

**Table 2:** Pathogenic profiles in known epilepsy genes. The 198 epileptic patients and 301 controls represent the discovery set. The replication set contains 325 epileptic patients and 380 controls. Variants that were not tested are marked with “-”.

| Patient | Epilepsy type | Syndrome | Copy number | Chr. | CNV start | CNV end | Epilepsy gene with exon disrupted | Taqman probe | Discovery |  | Replication |  |
| --- | --- | --- | --- | --- | --- | --- | --- | --- | --- | --- | --- | --- |
|  |  |  |  |  |  |  |  |  | Patients | Controls | Patients | Controls |
| CNET0108 | Generalized | Eyelid myoclonia epilepsy with absence | 1 | 1 | 44195001 | 44460000 | *ST3GAL3* | Hs05759463.cn | 1 DEL | 0 | 0 | - |
| CNET0159 | Generalized | Eyelid myoclonia epilepsy with absence | 1 | 8 | 141925001 | 142010000 | *PTK2* | Hs06202928.cn | 1 DEL | 0 | 0 | - |
| CNET0093 | Generalized | Juvenile onset; GTCs, Abs, Comp Partial | 1 | 10 | 95525001 | 95545000 | *LGI1* | Hs02682696.cn | 1 DEL | 0 | 0 | - |
| CNET0140 | Generalized | Idiopathic generalized epilepsies | 1 | 13 | 35750001 | 35785000 | *NBEA* | Hs05286691.cn | 1 DEL | 0 | 0 | - |
| CNET0144 | Generalized | Eyelid myoclonia epilepsy with absence | 1 | 15 | 22745001 | 23275000 | *NIPA2* | Hs04452887.cn | 3 DEL | 2 DEL | 4 DEL (2DUP) | 1 DEL (5 DUP) |
| CNET0009 | Generalized | Idiopathic generalized epilepsies | 1 | 15 | 30910001 | 32445000 | *CHRNA7* | Hs03909657.cn | 1 DEL | 0 | 3 DEL | (1 DUP) |
| CNET0119 | Generalized | Eyelid myoclonia epilepsy with absence | 1 | 15 | 93300001 | 93515000 | *CHD2* | Hs05385106.cn | 1 DEL | 0 | 0 | - |
| CNET0143 | Generalized | Childhood absence epilepsy | 1 |  | 93489776 | 93494317 |  | Hs026436998.cn | 1 DEL | 0 | 0 | - |
| CNET0130 | Generalized | Eyelid myoclonia epilepsy with absence | 3 |  | 93445001 | 93450000 |  | Hs01379802.cn | 1 DUP | 0 | 0 | - |
| CNET0074 | Focal | Frontal Lobe Epilepsy | 1 | 22 | 32125001 | 32255000 | *DEPDC5* | Hs01632214.cn | 1 DEL | 0 | 0 | - |

**Fig 4:**
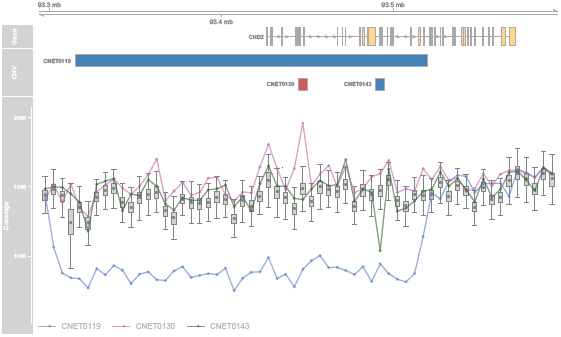
Exonic CNVs in *CHD2* detected by PopSV. The ‘CNV’ panel shows the exonic deletions (blue) and duplications (red) called by PopSV. The ‘Coverage’ panel shows the read depth signal in the affected individuals (colored points/lines) and the coverage distribution in the reference samples (boxplot and grey point).

Four of the 21 putative pathogenic CNVs were found in more than one individual (see Table 3 for precise numbers). To assess their global prevalence we tested them in an additional cohort of 325 epileptic patients and 380 ethnically matched controls (Table 3). Two regions were replicated:the first region in chromosome 2 consists of duplication of the genes *TTC27, LTPB1* and *BIRC6*. In total, 4 patients carried this duplication and it was not reported in any of the two sets of controls.

**Table 3:**
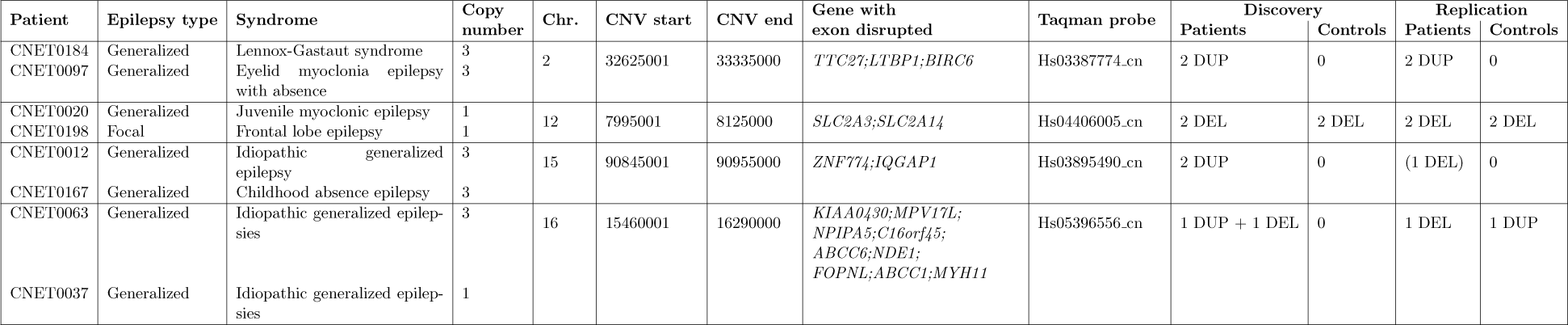
Recurrent CNVs with a pathogenic profile. The 198 epileptic patients and 301 controls represent the discovery set. The replication set contains 325 epileptic patients and 380 controls.

The second region was found on chromosome 16 and encompasses several genes. Two deletions were found in epileptic patients for this region and 1 epileptic individual and 1 control were also carriers of a duplication in the same region. This region corresponds to a genomic hotspot whose deletions were previously associated with epilepsy [30] and other neurological disorders. Finally, the remaining putative pathogenic CNVs were also associated with a number of genes (see Table S3). However, as we lack additional evidence for those specific CNV regions, we propose that these genes should be assessed in independent epilepsy cohorts. Of note, one patient had a rare 170 Kbp deletion encompassing three exons of the *PTPRD* gene which is predicted to be highly intolerant to loss-of-function mutations (pLI=1) [47]. Rare deletions in this gene were previously found in four independent individuals with attention-deficit hyperactivity disorder [55] and associated with intellectual disability [56]. In addition, de novo deletions were found in an individual with autism [57] and more recently in a patient with epileptic encephalopathy [32]. A common intronic variant in *PTPRD* was also associated with remission of seizures after treatment in a clinical cohort of epilepsy patients [58].

## 3 Discussion

Although several tools exist for the detection of CNVs using WGS data, we found that none of them could efficiently account for technical biases, thus resulting in limited sensitivity. To improve on this, we developed a new tool, PopSV, which we demonstrated was able to accurately detect CNVs, including rare and small events.

A key aspect of our approach is the use of a set of reference samples to identify abnormal read coverage. In this context, the choice and number of reference samples will have an effect on the analysis. Results from running PopSV using different reference cohort sizes suggest that CNV calls are consistent across runs but that a higher number of reference samples increases the sensitivity and robustness of the CNV detection (Fig S18). Based on these results, we recommend PopSV when 20 samples or more can be used as reference. In a given study, all samples can be used as a reference, or a subset of a few hundreds if the total sample size is extremely large. Although variants with frequency around 50% might not be detected, PopSV excels at detecting less frequent variants, smaller variants or variants in challenging regions such as repeat-rich regions. In a case/control design, the control samples could be used as reference in order to maximize the detection of case-specific variants. In the current study we used both epilepsy patients and controls as reference in order to be able to directly compare the observed CNV distributions. Finally, in a cancer project with paired normal and tumor samples, only normal samples should be used as reference such that PopSV can detect somatic CNVs of any frequency.

To maximize performance, the same library preparation, sequencing and data pre-processing should be employed on all the samples. To identify potential batch effects, a principal component analysis of read coverage was implemented as part of the PopSV package and is recommended to assess the homogeneity of the reference samples. The read length and aligner can lead to drastic changes in the read coverage and should be consistent across the cohort when analyzed with PopSV. This is particularly important in repeat-rich regions. Although the different datasets were produced by different sequencing and pre-processing protocols and showed varying degrees of technical bias (Fig 1a, S1 and S2), the performance of PopSV was comparable when benchmarking the methods in the two public datasets and experimentally validating calls in the CENet cohort.

PopSV’s approach does not require a uniform read coverage and integrate the coverage variation separately in each studied region. For these reasons, it would be straightforward to analyze targeted sequencing data, such as exome-sequencing. PopSV could also be extended for the detection of other types of SVs such as balanced SVs. To do this, instead of counting properly mapped reads, the method could be modified to test for an excess of discordant reads. Finally, additional modules could be added to PopSV to help characterize the detected variants. For instance, instead of computing a copy-number estimate from the average coverage in the reference, a HMM approach including all samples could provide a better genotyping strategy. Similar to other approaches [9,16], an additional step in the pipeline could explore the effect of the bin size on the variation in read coverage across the population and suggest an optimal bin size.

As in previous array-based studies [35–37], we observed an enrichment of large rare exonic CNVs in patients compared to controls. However, thanks to the resolution of WGS and PopSV, we found that the global distribution of small CNVs (*<*50 Kbp) in 198 unrelated epilepsy patients was also skewed towards rare exonic CNVs. In addition, genes disrupted by rare deletions in patients were enriched for previously known epilepsy genes. These observations support the association of small CNVs with epilepsy and could not have been detected in previous array-based studies.

We also observed a clear enrichment of non-coding CNVs in the neighborhood of previously implicated genes. When focusing on CNVs seen only in the epilepsy cohort and around epilepsy genes, 10.1% of epilepsy patients have an exonic CNVs and our results shows that up to 28.8% of patients harbor non-coding CNVs of high-interest in the proximity of epilepsy genes. These non-coding variants are present in the epilepsy cohort only and located in annotated regulatory regions associated to known epilepsy genes. Although it is challenging to directly test their functional impact, their frequency and location suggest a putative importance in the genetic mechanism of epilepsy and should be further investigated in the future.

Finally, to better understand the impact of these findings on an individual scale, we selected CNVs with the highest pathogenic potential within our patients. These CNVs highlighted known but also potentially new epilepsy genes. Using a second epilepsy cohort, we were also able to identify two chromosomal regions that were recurrently disrupted by CNVs. These findings highlight the benefits of having a comprehensive survey of CNVs when trying to understand the genetic causes of a disease.

## 4 Materials and methods

### 4.1 Epilepsy patients and sequencing

Patients were recruited through two main recruitment sites at the Centre Hospitalier Universitaire de Montréal (CHUM) and the Sick Kids Hospital in Toronto as part of the Canadian Epilepsy Network (CENet). This study was approved by the Research Ethics Board at the Sick Kids Hospital (REB number `1000033784`) and the ethics committee at the Centre Hospitalier Universitaire de Montréal (project number `2003-1394,ND02.058-BSP(CA)`). Before their inclusion in this study, patients or parents (when needed) had to give written informed consents. The main cohort of this study was constituted of 198 unrelated patients with various types of epilepsy; 85 males and 113 females. The mean age at onset of the disease for our cohort was 9.2 (*±*6.7) years. Supplementary Table S1 presents a detailed description of the clinical features for the various individuals recruited in this study. 301 unrelated healthy parents of other probands from CENet were also included in this study and used as a control cohort. DNA was exclusively extracted from blood DNA.

Libraries were generated using the TruSeq DNA PCR-Free Library Preparation Kit (Illumina) and paired-end reads of size 125 bp were sequenced on a HiSeq 2500 to an average coverage of 37.6x *±* 5.6x. Reads were aligned to reference Homo sapiens b37 with BWA [59]. Finally, Picard was used to merge, realign and mark duplicate reads. Raw sequence data has been deposited in the European Genome-phenome Archive, under the accession code EGAS00001002825. For more details, see Supplementary Information.

### 4.2 Public WGS datasets

Two high-coverage public datasets were used to benchmark PopSV against existing methods.

A *Twin* study provided WGS sequencing data for 45 individuals, including 10 monozygotic twin quartets from the Quebec Study of Newborn Twins [38]. All patients gave informed consent in written form to participate in the Quebec Study of Newborn Twins. Ethic boards from the Centre de Recherche du CHUM, from the Université Laval and from the Montreal Neurological Institute approved this study. DNA was extracted from blood and sequencing was done on an Illumina HiSeq 2500 (paired-end mode, fragment length 300 bp). The reads were aligned using a modified version of the Burrows-Wheeler Aligner [59] (bwa version 0.6.2-r126-tpx with threading enabled). The options were `’bwa aln -t 12 -q 5’ and ‘bwa sampe -t 12’`. Aligned reads are available on the European Nucleotide Archive under ENA PRJEB8308. The 45 samples had an average sequencing depth of 40x (minimum 34x / maximum 57x).

A cancer dataset from a study of renal cell carcinoma [39] was also used. 95 pairs of normal/tumor tissues were sequenced using GAIIx and HiSeq2000 instruments. Paired-end reads of size 100 bp totaled an average sequencing depth of 54x (minimum 26x / maximum 164x). Reads were trimmed with FASTX-Toolkit and mapped per lane with BWA [59] backtrack to the GRCh37 reference genome. Picard was used to adjust pairs coordinates, flag duplicates and merge lanes. Finally, realignment was done with GATK. Raw sequence data has been deposited in the European Genome-phenome Archive, under the accession code EGAS00001000083. More details can be found in Scelo et al. [39].

### 4.3 Testing for technical biases in WGS

To investigate the bias in read depth (RD), we fragmented the genome in non-overlapping bins of 5 Kbp and counted the number of properly mapped reads. In each sample, we corrected for GC bias and removed bins with extremely low or high coverage (see Supplementary Information). Then, read counts across all samples were combined and quantile-normalized. Using simulations and permutations, we constructed two control RD datasets with no region-specific or sample-specific bias. We computed the mean and standard deviation of the coverage in each bin across samples. Next, to investigate experiment-specific bias, we retrieved which sample had the highest coverage in each bin. Then we computed, for each sample, the proportion of the genome where it had the highest coverage. The same analysis was performed monitoring the lowest coverage. This analysis was performed separately on the CENet dataset, the Twin dataset and the normal samples from the cancer dataset. On the Twin dataset, the same analysis was also run after correcting the read coverage following the QDNAseq pipeline [40] (see Supplementary Information).

### 4.4 PopSV

The main idea behind PopSV is to assess whether the coverage observed in a given location of the genome diverges significantly from the coverage observed in a set of reference samples. PopSV was implemented in an R package (see Data and code availability). The genome is first segmented into bins and the number of reads with proper mapping in each bin is counted for each sample. In a typical design, the genome is segmented in non-overlapping consecutive windows of equal size, but custom designs could also be used. With PopSV, we propose a new normalization procedure which we call targeted normalization that retrieves, for each bin, other genomic regions with similar profile across the reference samples and uses these bins to normalize read coverage (see Supplementary Information). Our targeted normalization was compared to global approaches that adjust for the median coverage, or quantile-based approaches. After normalization, the value observed in each bin is compared with the profiles observed in the reference samples and a Z-score is calculated (Fig 1b). False Discovery Rate (FDR) is estimated based on these Z-score distributions and a bin is marked as abnormal based on a user-defined FDR threshold. Consecutive abnormal bins are merged and considered as one variant. In PopSV’s R package, circular binary segmentation [60] can also be used to merge bins into variant regions. Copy number was estimated by dividing the coverage in a region by the average coverage across the reference samples, multiplied by 2 (see Supplementary Information).

### 4.5 Validation and benchmark of PopSV

We compared PopSV to CNVnator [9], FREEC [10] and cn.MOPS [11], three popular RD methods that can be applied to WGS datasets. We also ran LUMPY [8] which uses an orthogonal mapping signal: the insert size, orientation and split mapping of paired reads. For LUMPY, all the CNVs (deletions and duplications) and intra-chromosomal translocations (labeled as ‘BND’ in Lumpy’s output) larger than 300 bp were kept for the upcoming analysis. These methods were run on the two publicly available datasets, using 5 Kbp bins for the RD methods.

First, we compared the frequency at which a region is affected by a CNV using the calls from the different methods. To investigate the presence of systematic calls in each method, we compute how many of the calls in a typical sample are called at different frequencies in the dataset. For example, on average, how many calls in one sample are called in more than 90% of the samples. In the Twin dataset, the samples were clustered using the CNV calls from each method. Different linkage criteria were used for the hierarchical clustering (see Supplementary Information). The Rand index estimated the concordance between the clustering and the known pedigree (family-level). Next, we measured the number of CNVs identified in each twin that were also found in their monozygotic twin. We removed calls present in more than 50% of the samples to ensure that systematic errors were not biasing our replication estimates. Hence, a replicated call is most likely true as it is present in a minority of samples but consistently in the twin pair. For CNVnator, LUMPY and PopSV, the eval1/eval2 columns, number of supporting reads and adjusted P-values (respectively) were used to gradually filter low-quality calls and explore their effect on the replication metrics. In addition to their replication, we annotated the calls when their region overlapped a call found by other methods in the same sample. For calls found by at least two methods, we computed the proportion of calls from a method found by each of the other methods.

The approach described previously comparing pairs of twins was also applied in the cancer dataset, on pairs of normal/tumor samples. In this case, a replicated call is found in the normal sample and in the paired tumor sample. Finally, we compared calls using small bins (500 bp) and calls using larger bins (5 Kbp). This comparison explores the quality of the calls, the size of detectable events and the resolution for different bin sizes. First, we counted how many small bin calls supported any large bin call. We then looked at the proportion of small bin calls of different sizes that were also found in the large bin calls.

### 4.6 CNV detection in the CENet cohorts

CNVs were called using PopSV using 5 Kbp bins and all the samples from both the epilepsy and control cohorts as reference. We annotated the frequency of the CNVs using germline CNV calls from the Twin and cancer datasets (internal database) as well as four public CNV databases from the 1000 Genomes Project [13, 45], the Genome of Netherlands [44] and the Simons Genome Diversity Project [46]. CNVs were annotated with the maximum frequency in the databases. Hence, a rare CNV is defined as present in less than 1% of the samples in each of the five CNV databases.

To test for a difference in deletion/duplication ratio among rare CNVs, we compared the numbers of rare deletions and duplications in the epilepsy patients and controls using a *χ*^2^ test. The same test was performed after downsampling the controls to the sample size of the epilepsy cohort.

### 4.7 Validation by Taqman RT-PCR

We first selected CNV calls in epilepsy patients that spanned at least 2 consecutive bins. We kept exonic CNVs of different sizes and overlapping a Taqman probe. A second batch of CNVs, containing small non-coding CNVs, was also sent for validation. Here, hundreds of non-coding CNVs spanning only one bin were randomly selected. When possible the breakpoints were manually fine-tuned from manual inspection of a base-pair level coverage representation or using IGV [61]; the breakpoints remained unchanged when they could not be refined. Finally, we kept regions overlapping a Taqman probe.

Probes were selected using the assay search tool on the Thermofisher website. All probes were tested for patients and controls that were called in PopSV as well as an additional 10 control individuals to ensure the validity of the probe. For each CNV, one assay was chosen in the middle of the genomic region of interest and located in an exon when possible. All reactions with TaqMan Copy Number Assays were performed in duplex using the FAM dye label based assay for the target of interest (Taqman copy number assay, Made to order, #4400291, Applied Biosystems by Life Technologies) and the VIC dye label based TaqMan Copy Number Reference Assay for RNase P (4403326, Life technologies). Amplification reactions (10*µL*), which were performed in quadruplicate, consisted of: 10 ng gDNA, 1X TaqMan Copy Number Assay, 1X TaqMan Copy Number Reference Assay, RNase P, 1X TaqMan Genotyping Master Mix (4371355, Life Technologies) or 1X SensiFAST Probe Lo-ROX Kit (BIO-84020, Froggabio). PCR was performed with an Applied Biosystems QuantStudio7 flex Real-Time PCR system using the standard curve settings and the default universal cycling conditions: 95 °C 10 minutes followed by 40 cycles: 95 °C 15 seconds, 60 °C 60 seconds. Data was analyzed with QuantStudio Real-Time PCR system software v1.2 (Applied Biosystems by Life Technologies) using autobaseline and manual Ct threshold of 0.2. Results export files were opened in CopyCaller^TM^ Software v2.0 for sample copy number analysis by the relative quantitation method. The median ΔCt was used as the calibrator sample in the analysis settings.

### 4.8 CNV enrichment in exonic regions

For each cohort (epilepsy and control), we retrieved the CNV catalog by merging CNV that are recurrent in multiple samples. Hence, the CNV catalog represents all the different CNVs found in each cohort. Because the epilepsy and control cohorts have different sample sizes, the CNV catalogs for each cohort were built using 150 randomly selected samples. For each sub-sampling and each cohort, control regions were selected to fit the size distribution of the CNV catalog and the overlap with centromeres, telomeres and assembly gaps (see Supplementary Information). The fold-enrichment represents how much more/less of the CNVs overlap an exon compared to the control regions. To robustly compare the two cohorts, we computed the median difference in fold-enrichment between the CNV catalogs from patients and controls across 100 sub-sampled catalogs. The cohort labels of the CNV catalogs were then permuted 10,000 times and the analysis repeated to derive a null distribution for the median difference in fold enrichment. A permuted P-value was computed from the observed difference and the null distribution.

Small (*<*50Kbp) and large (*>*50 Kbp) CNVs were analyzed separately. Exons from genes predicted to be loss-of-function intolerant [47] (probability of loss-of-function intolerance *>* 0.9) were also analyzed separately. The same analysis was repeated using only rare CNVs, i.e. being present in less than 1% of PopSV calls in the Twins and renal cancer datasets, and in four public datasets (see Supplementary Information).

In each cohort, we then retrieved the CNV catalog of rare exonic CNVs. We evaluated the proportion of the CNVs in the catalog that are private (i.e. seen in only one sample). The control cohort was down-sampled a thousand times to the same sample size as the epilepsy cohort to provide a confidence interval and empirical P-value (see Supplementary Information). We also visualize the proportion of CNVs in the catalog seen in 2 samples or more, 3 samples or more, etc (Fig S12a). We performed the same analysis after removing the top 20 samples with the highest number of non-private rare exonic CNVs. The analysis was also repeated using French-Canadian individuals only.

### 4.9 CNV enrichment in and near epilepsy genes

We used the list of genes associated with epilepsy from the EpilepsyGene resource [48] which consists of 154 genes strongly associated with epilepsy. We tested different sets of CNVs: deletion or duplications in the epilepsy cohort, control individuals and samples from the twin study, and using different threshold of maximum frequency. For each set of CNVs, we counted how many of the genes hit were known epilepsy genes. To control for the size of epilepsy genes and CNV-hit genes, we randomly selected genes with sizes similar to the genes hit by CNVs and evaluated how many were epilepsy genes. After sampling 10,000 gene sets, we computed an empirical P-value (see Supplementary Information).

To investigate rare non-coding CNVs close to known epilepsy genes, we counted how many patients have such a CNV at different thresholds of distance to the nearest exon. We compared this cumulative distribution to the control cohort, after down-sampling it to the sample size as the epilepsy cohort. We performed the same analysis using deletions only. Each epilepsy gene was also tested for an excess of rare non-coding deletions in patients versus controls using a Fisher test. Next, we restricted our analysis to rare non-coding CNVs that overlap an eQTL associated with the epilepsy genes [50] or a DNase I hypersensitive site associated with the promoter of epilepsy genes [51]. A Kolmogorov-Smirnov test was used to test the difference in distribution. Finally, using different values for the maximum distance to the nearest epilepsy gene, we computed the odds ratio of having such a CNV between epilepsy patients and controls.

### 4.10 Putatively pathogenic CNVs

Exonic CNVs larger than 10 Kbp and found in less than 1% of the 301 controls were first selected. We further retained either CNVs overlapping the exon of a known epilepsy-associated gene [48] or deletions overlapping the exon of a loss-of-function intolerant gene [47], or CNVs present in two or more of our epilepsy patients. All the putatively pathogenic CNVs were validated by Taqman RT-PCR.

### 4.11 Data and code availability

The PopSV R package and its documentation are available at http://jmonlong.github.io/PopSV/. Scripts are provided to run the pipeline on different high performance computing systems. The code used for the analysis and to produce figures and numbers is documented at http://github.com/jmonlong/epipopsv and archived in https://doi.org/10.5281/zenodo. 1172181. Necessary data, including the CNV calls, was deposited at https://figshare.com/s/20dfdedcc4718e465185. Raw sequence data has been deposited in the European Genome-phenome Archive, under the accession code EGAS00001002825.

## 5 Acknowledgments

Data analyses were enabled by compute and storage resources provided by Compute Canada and Calcul Québec. We would like to thank Pascale Marquis at the Canadian Centre for Computational Genomics for processing the raw sequencing data to genomic variant calls and for her active participation in various quality assessment steps. The Canadian Centre for Computational Genomics (C3G) is a Node of the Canadian Genomic Innovation Network and is supported by the Canadian Government through Genome Canada. We are grateful to the team of the Québec Study of Newborn Twins who provided the twin dataset and the Cagekid consortium who provided the renal cancer dataset. We would like to thank Sylvia Dobrzeniacka for sample handling and lab work. We are grateful to Dr. Ledia Brunga for her work on the epileptic cohort and to Brianna Goldenstein and Claudia Moreau for revising this manuscript. Finally, we would like to thank Simon Gravel, Mathieu Blanchette, Mathieu Bourgey, Toby Dylan Hocking and Claudia Moreau for helpful discussions.

## 6 Supplementary Tables

Table S1: **Clinical features of epileptic patients.** The Excel file contains the type of epilepsy, age of onset, sex, family history, pharmaco-resistance and potential intellectual disabilities.

Table S2: **PopSV calls validated by RT-PCR.** The Excel file contains the location of each region, the CNV type, the number of carriers in the CENet cohorts, the maximum proportion of carriers in the CNV databases, Taqman probe ID and validation status.

**Table S3:**
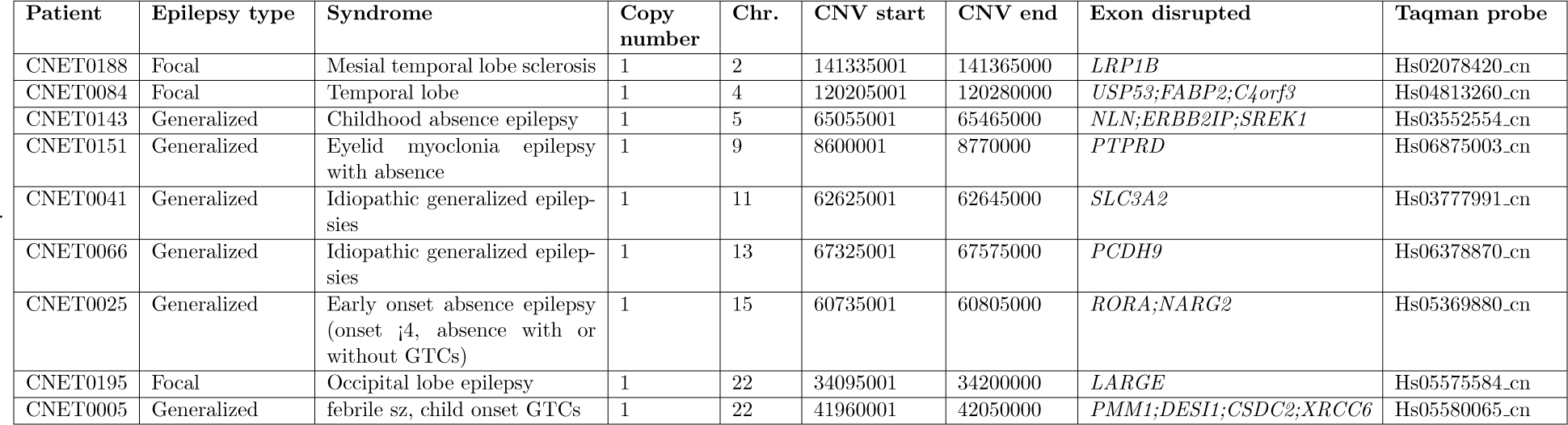
Other pathogenic profiles.

## 7 Supplementary Figures

**Fig S1:**
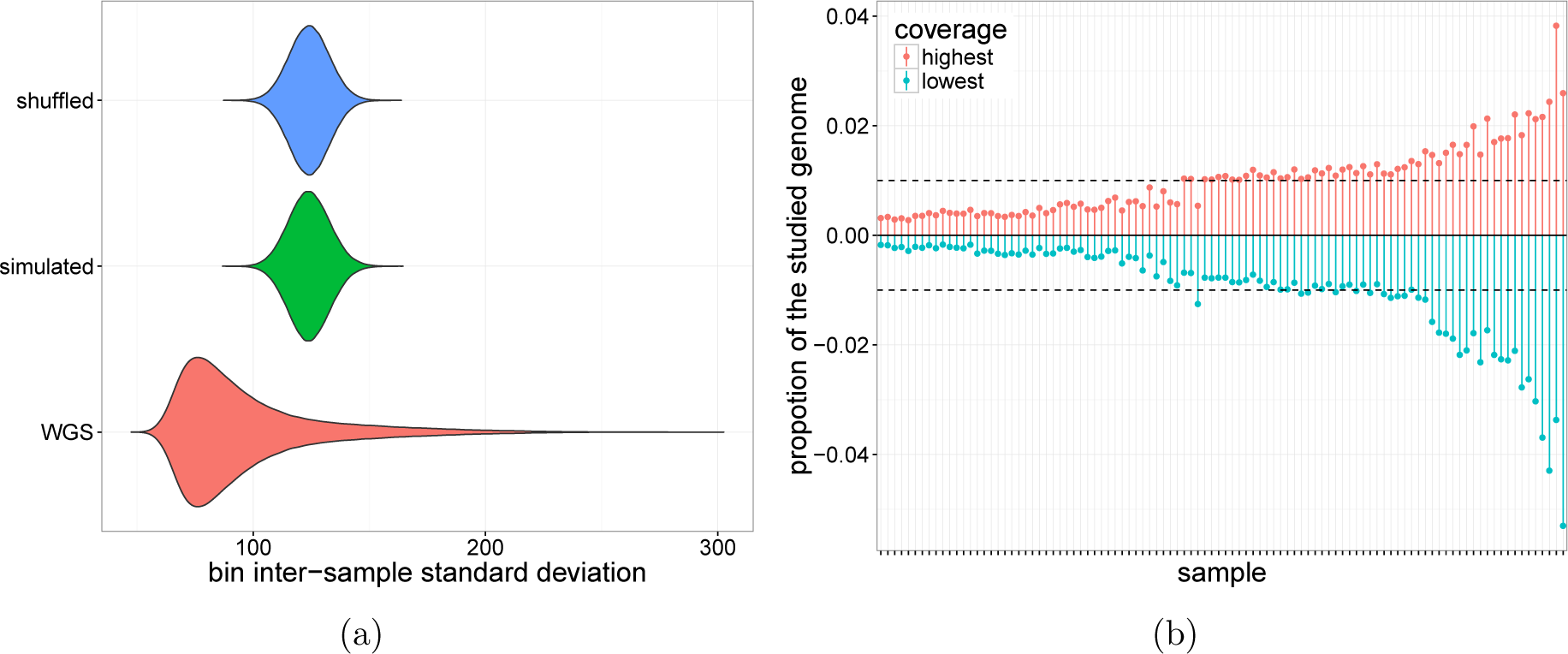
Variation and bias in whole-genome sequencing experiments in the epilepsy cohort. a) Distribution of the bin inter-sample standard deviation coverage (red) and null distribution (blue: bins shuffled, green: simulated normal distribution). b) Proportion of the genome in which a given sample (x-axis) has the highest (red) or lowest (blue) RD. In the absence of bias all samples should be the most extreme at the same frequency (dotted horizontal line).

**Fig S2:**
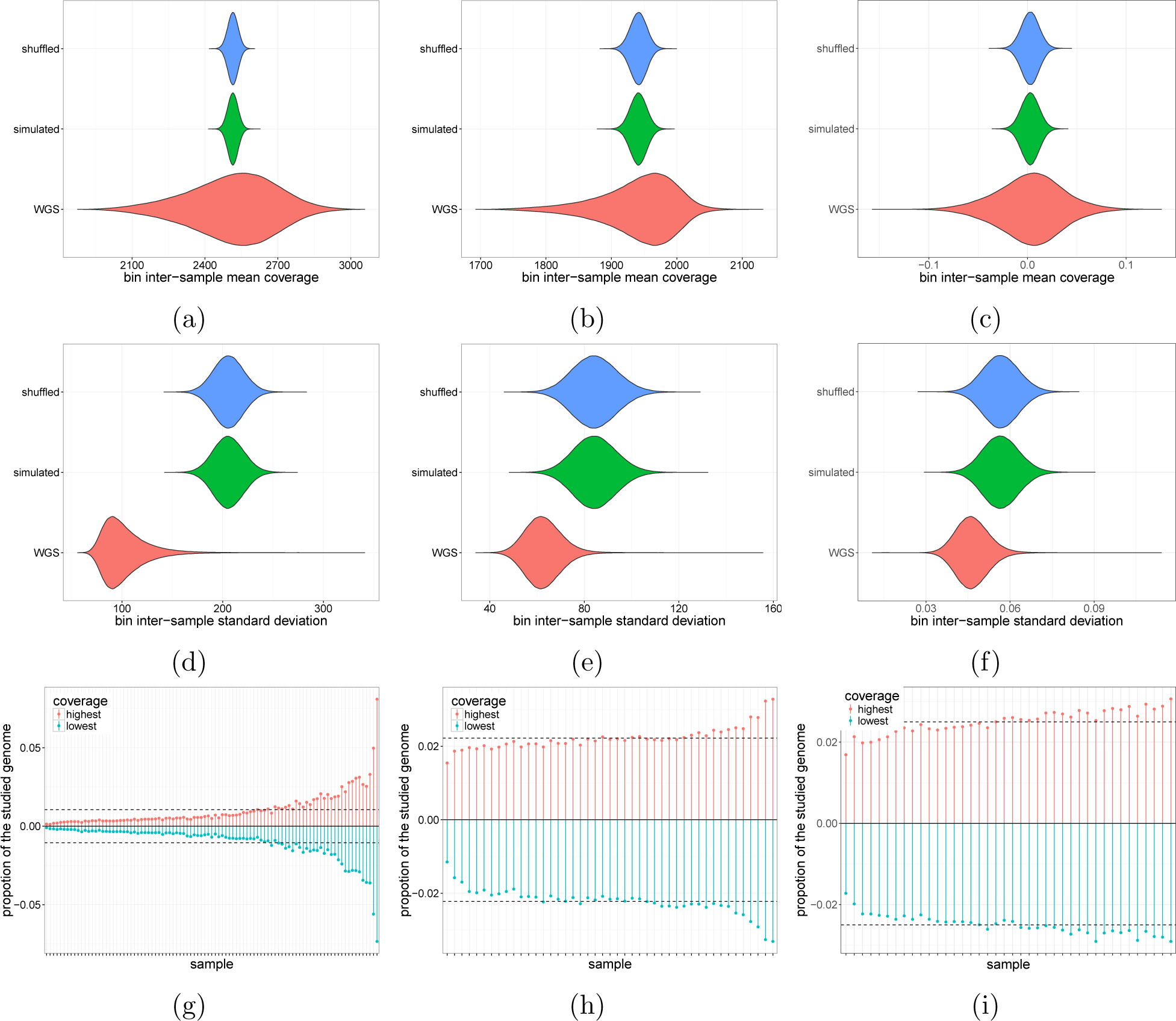
Variation and bias in whole-genome sequencing experiments in the normals from CageKid (a,d,g), the twin dataset (b,e,h) and the twin dataset after using QD- NAseq [40] correction (c,f,i). a-c) Distribution of the bin inter-sample standard deviation coverage (red) and null distribution (blue: bins shuffled, green: simulated normal distribution). d-f) Same for the bin inter-sample standard deviation coverage. g-i) Proportion of the genome in which a given sample (x-axis) has the highest (red) or lowest (blue) RD. In the absence of bias all samples should be the most extreme at the same frequency (dotted horizontal line).

**Fig S3:**
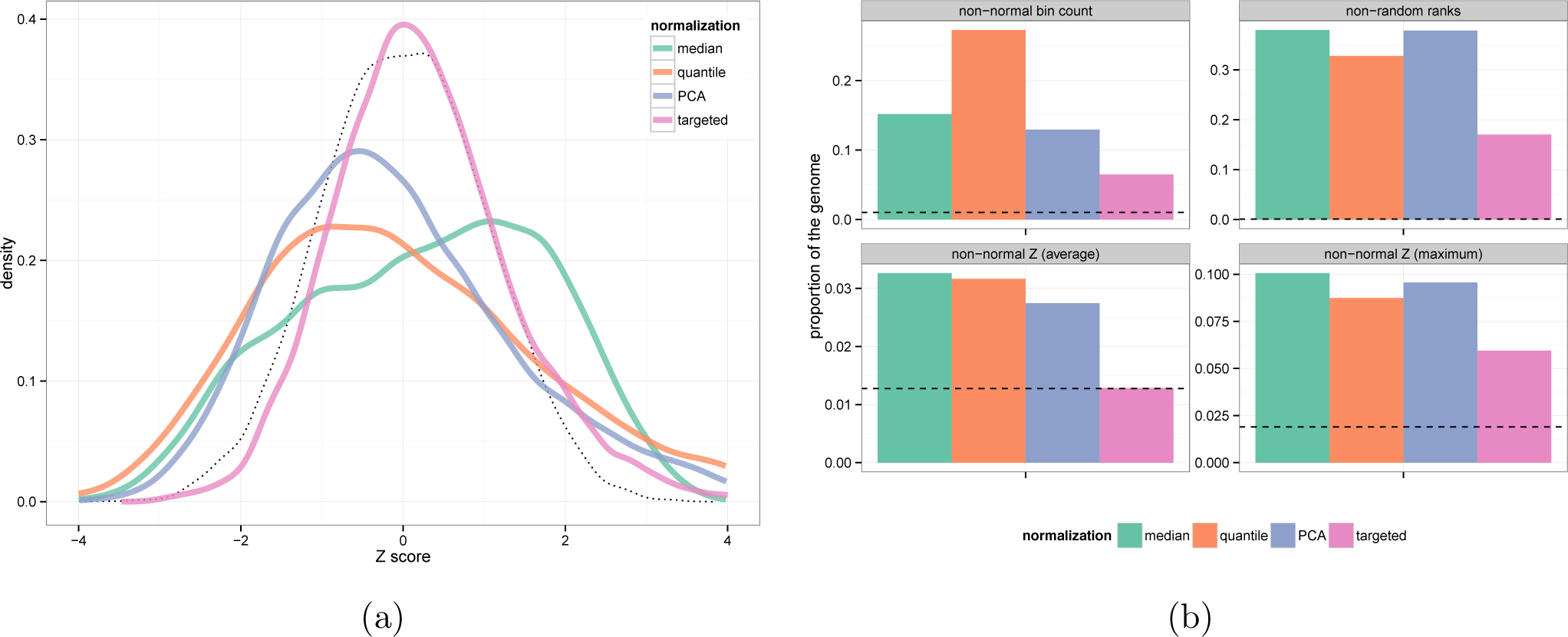
Comparison of different normalization approaches. a) For each normalization approach, the sample with the least normal Z-score distribution is shown. b) After targeted normalization, a lower proportion of the genome looks problematic for the analysis. Fewer bins have non-normal bin counts (top-left), the sample ranks are more random suggesting less sample-specific bias (top-right), and Z-scores fit better a Normal distribution on average (bottom-left) and in the worst sample (bottom-right). The dotted line is computed from simulated bin counts.

**Fig S4:**
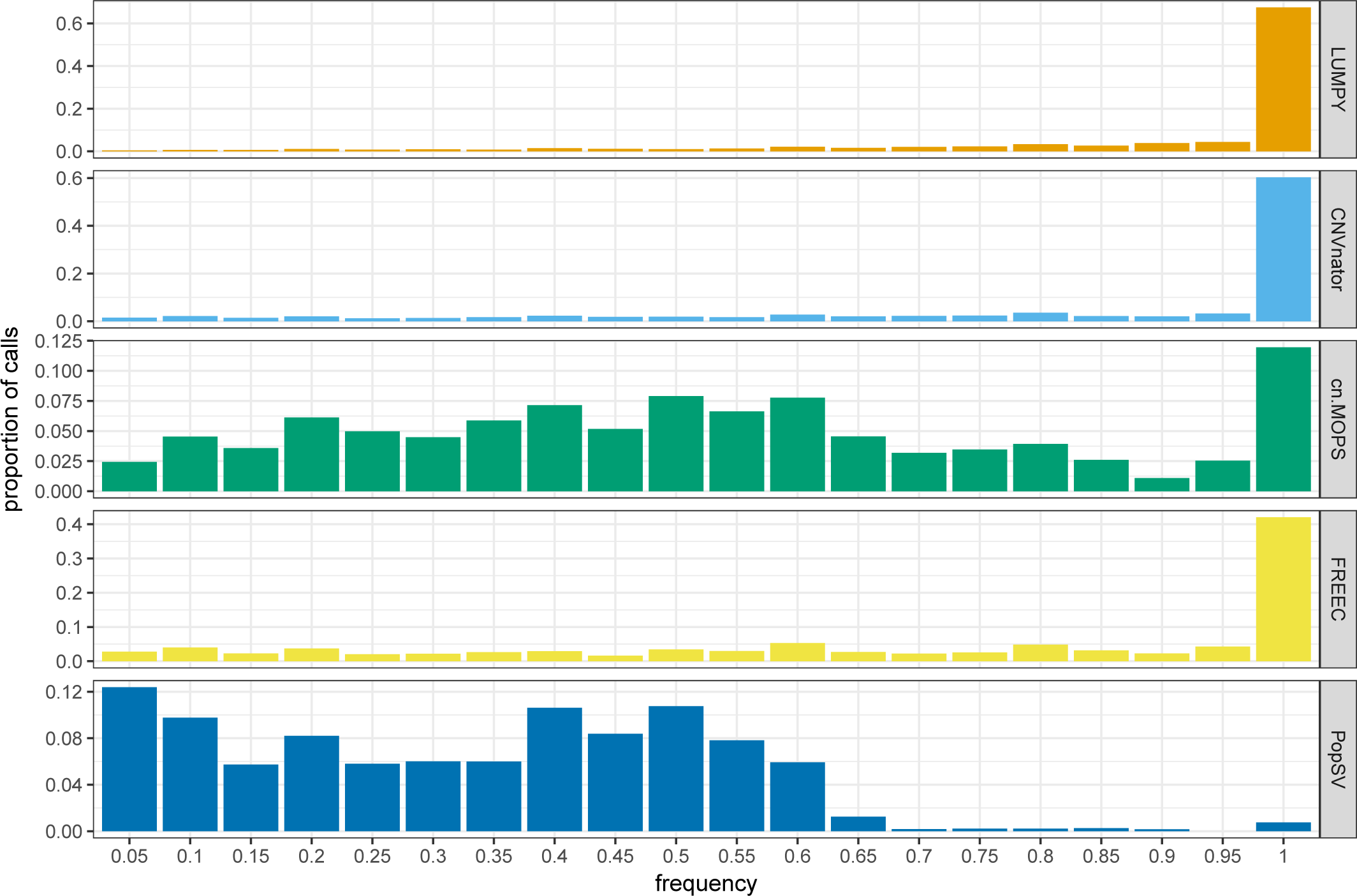
Frequency of calls in an average sample from the Twin study. The bars show the proportion of calls in an average samples (y-axis), grouped by the frequency of the call in the dataset (x-axis), for different methods.

**Fig S5:**
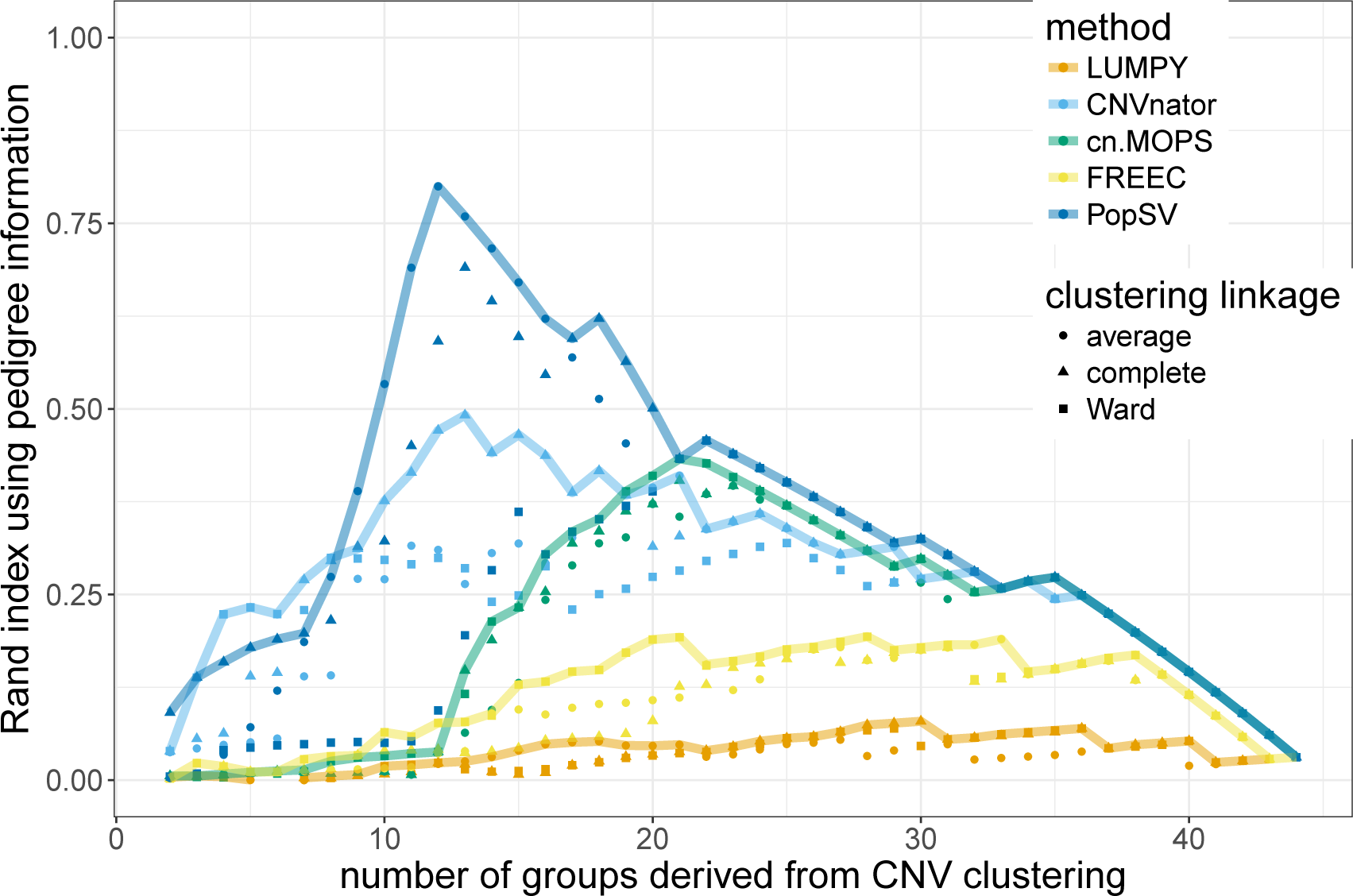
CNV clustering and twin pedigree. The hierarchical cluster tree from the CNV calls is cut at different levels (*x-axis*), cluster groups are compared to the known pedigree using the Rand index (*y-axis*). Different clustering linkage criteria (*point style*) are used and the one showing the best Rand index is highlighted by the line

**Fig S6:**
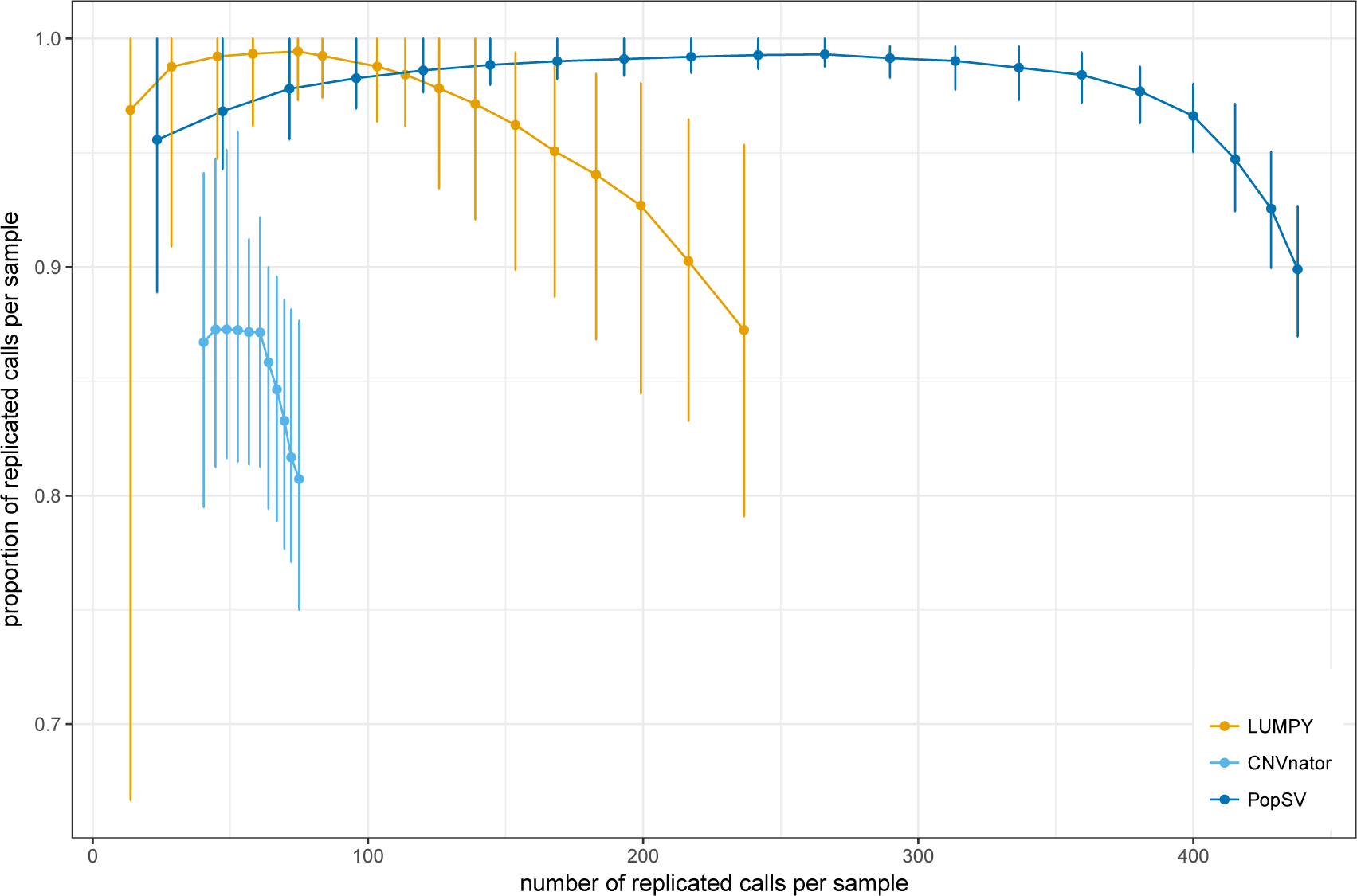
Replication in twins for different significance thresholds. Each point represents the number of replicated calls per sample (average across samples) and the proportion of replicated calls per sample. The vertical error bar shows the variation of the replication rate across the samples. The points and lines were computed by filtering calls at different significance levels (q-value for PopSV, number of supporting reads for LUMPY and eval1/eval2 for CNVnator, see Supplementary Information).

**Fig S7:**
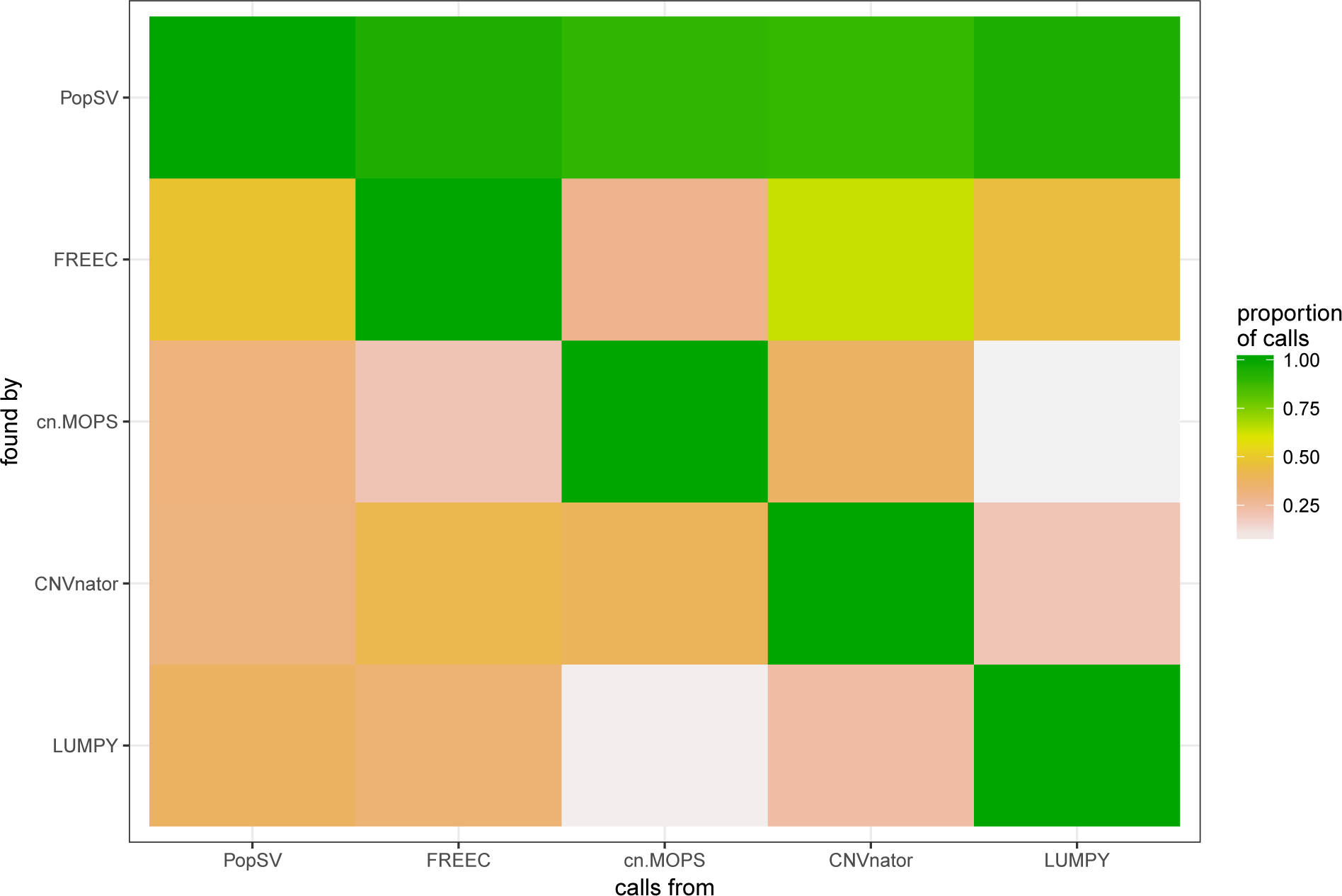
Calls found by several methods. Focusing on calls found by at least two methods, the heatmap shows the proportion of calls from one method (x-axis) that were also found by another (y-axis) on average per sample.

**Fig S8:**
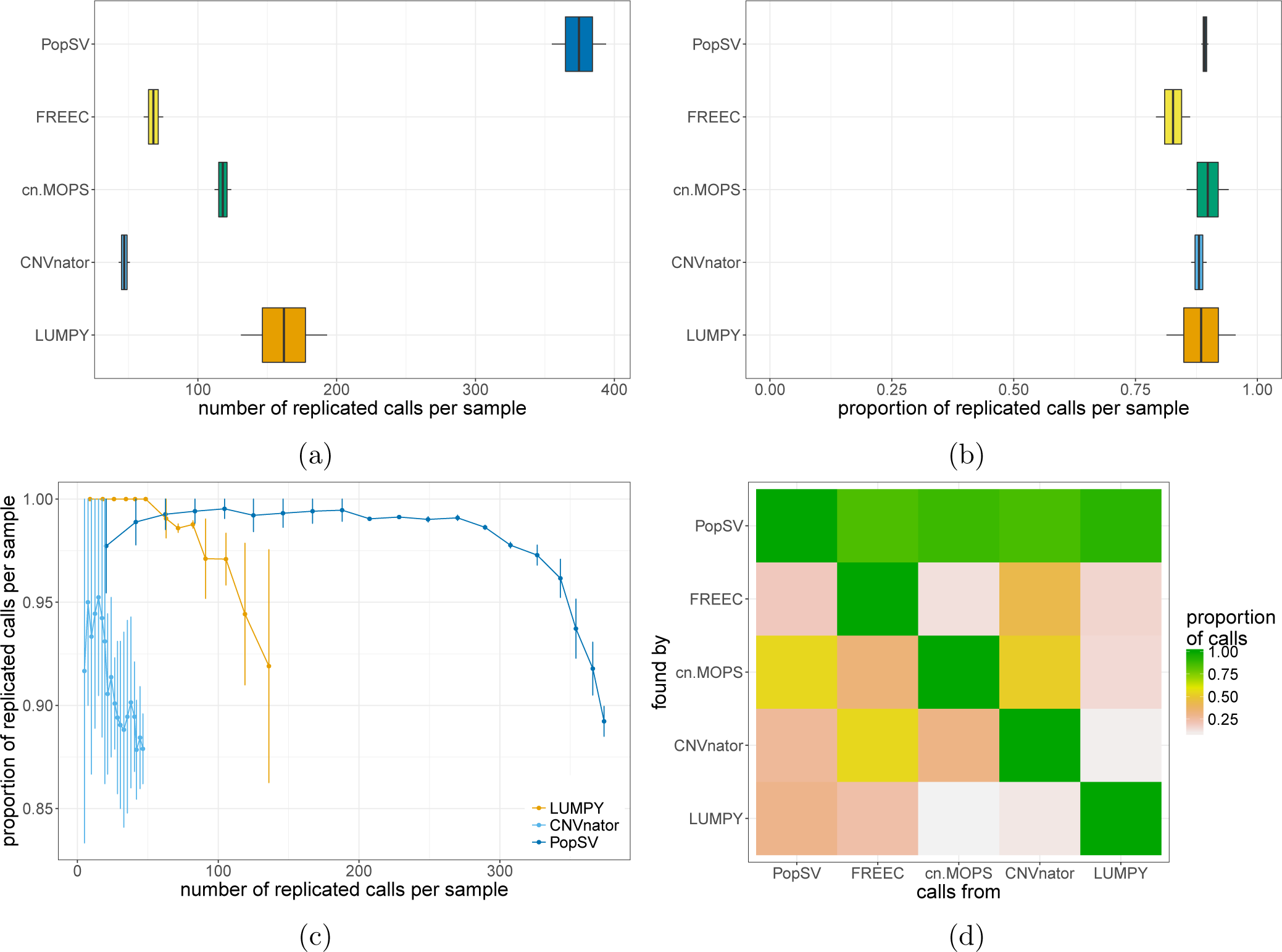
Benchmark across paired normal/tumor in CageKid. Number (a) and proportion (b) of germline calls replicated in the paired tumor in CageKid. c) Number and proportion of replicated calls when filtering calls at different significance levels. d) Focusing on calls found by at least two methods, the color shows the proportion of calls from one method (x-axis) that were also found by another (y-axis) on average per sample.

**Fig S9:**
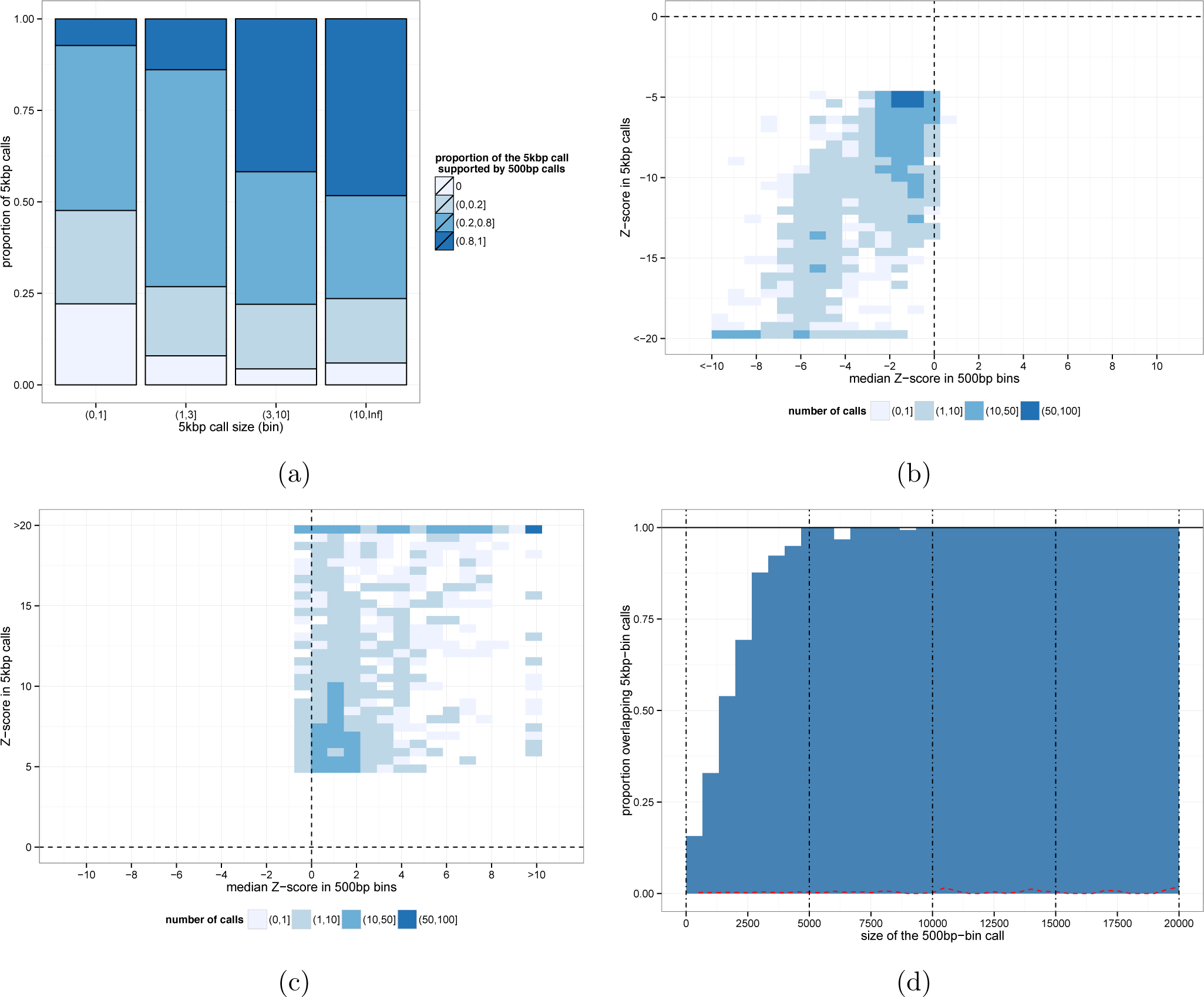
Comparison of PopSV results using different bin sizes. a) 5 Kbp calls of different sizes (x-axis) are split according to the proportion of the call supported by 500 bp calls. The Z-score of 500 bp bins in 5 Kbp calls is consistent with the call for deletion b) and duplication c) signal. 5 Kbp calls with lower significance (e.g. single-bin calls) are less supported by 500 bp calls (a) but their Z-scores are in the consistent direction (b,c) although not always significant enough to be called. d) Proportion of 500 bp calls of different sizes (x-axis) overlapping a 5 Kbp call.

**Fig S10:**
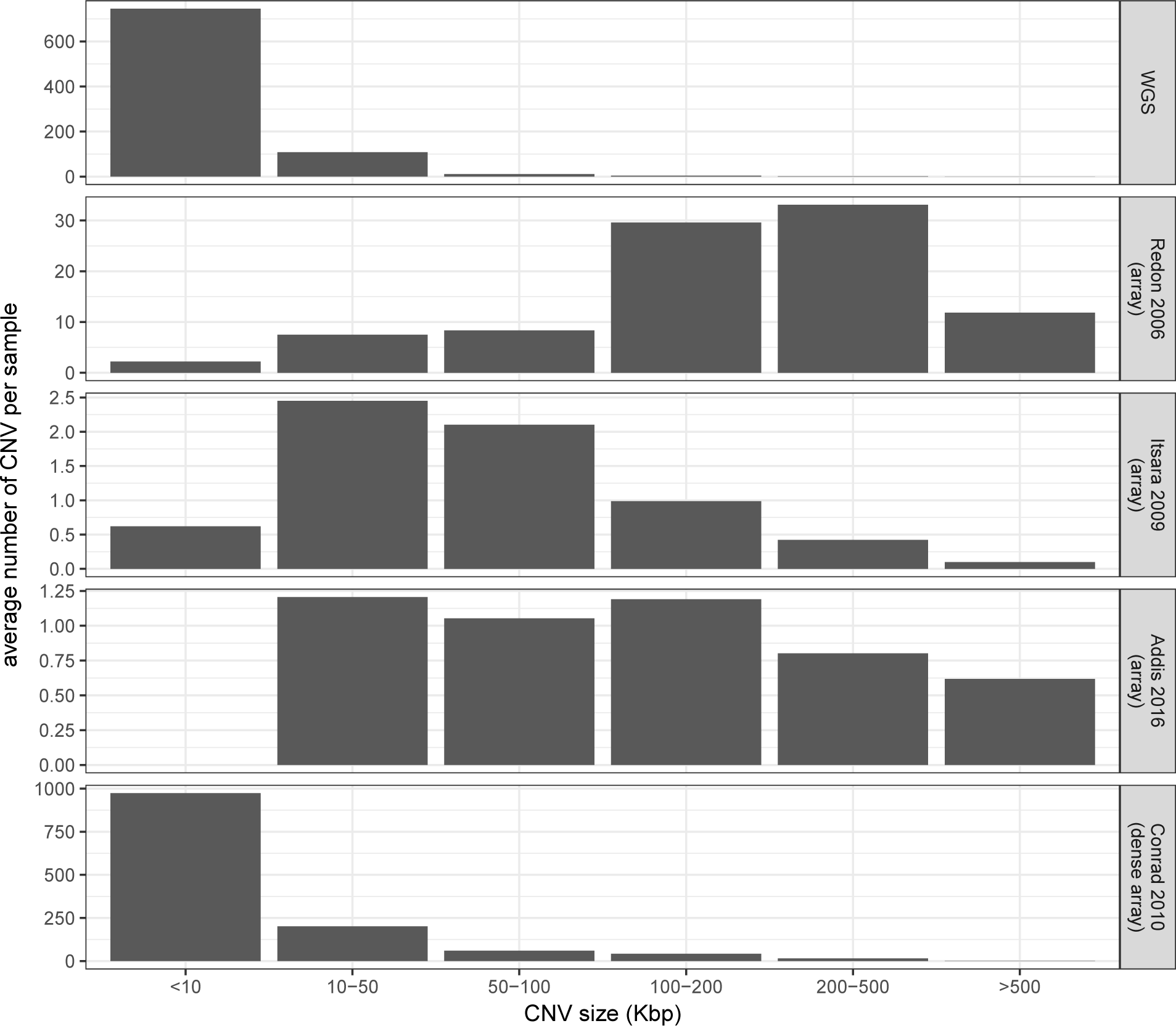
CNV size in our cohort and four array-based studies. The bars show the average number of CNVs called in a sample in the different cohorts. *Redon 2006* [42] and *Itsara 2009* [43] are population studies using technology similar to previous epilepsy studies. *Addis 2016* [34] is a recent study of large CNVs in absence epilepsy. *Conrad 2010* [4] is a population study that used multiple arrays to increase its resolution.

**Fig S11:**
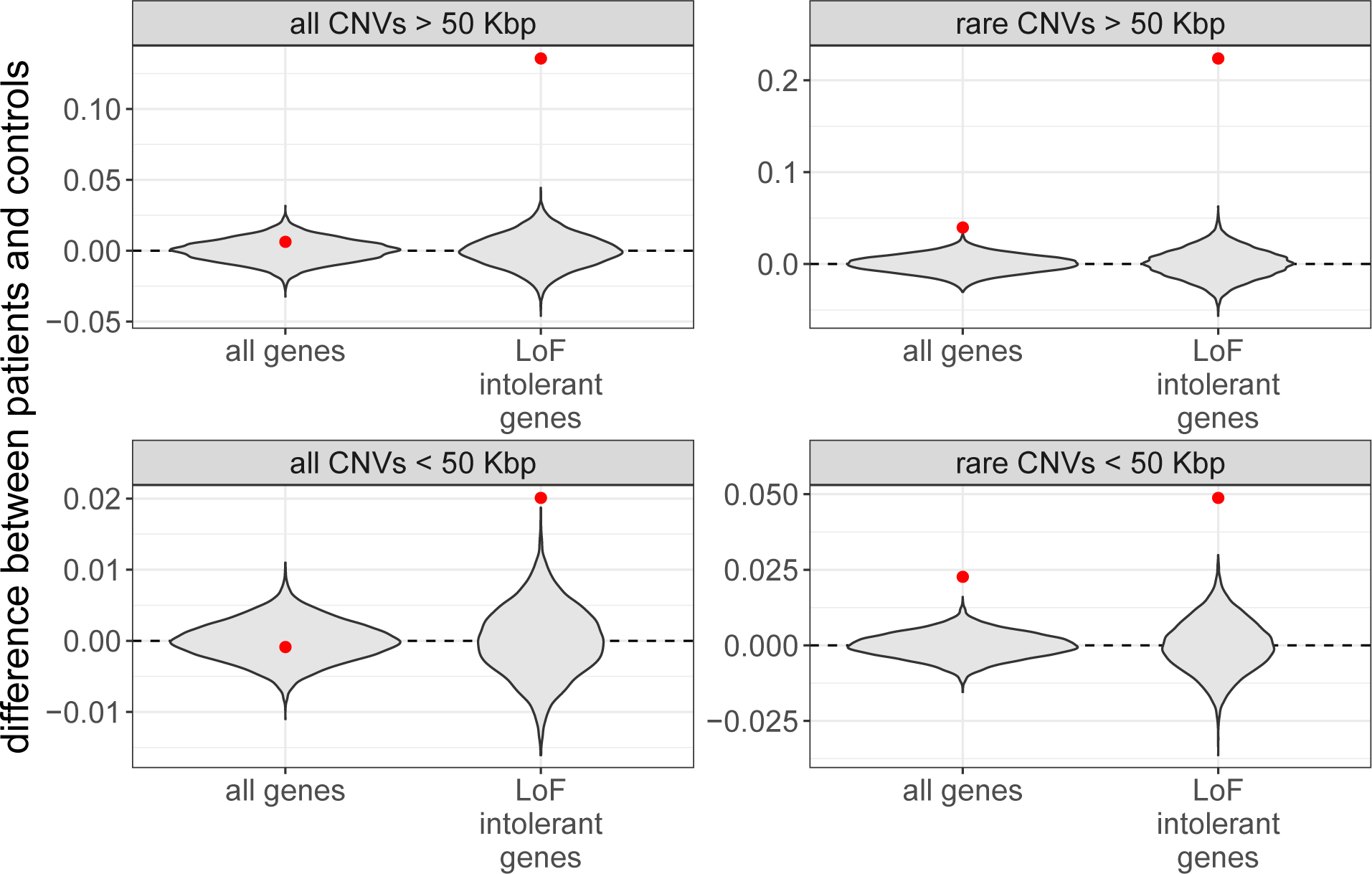
Exonic enrichment significance. The grey violin plot represents the difference in fold-enrichment between patients and controls across 10,000 permutations where the patient/control labels had been shuffled. The red point represents the observed difference between patients and controls (Figure 2c).

**Fig S12:**
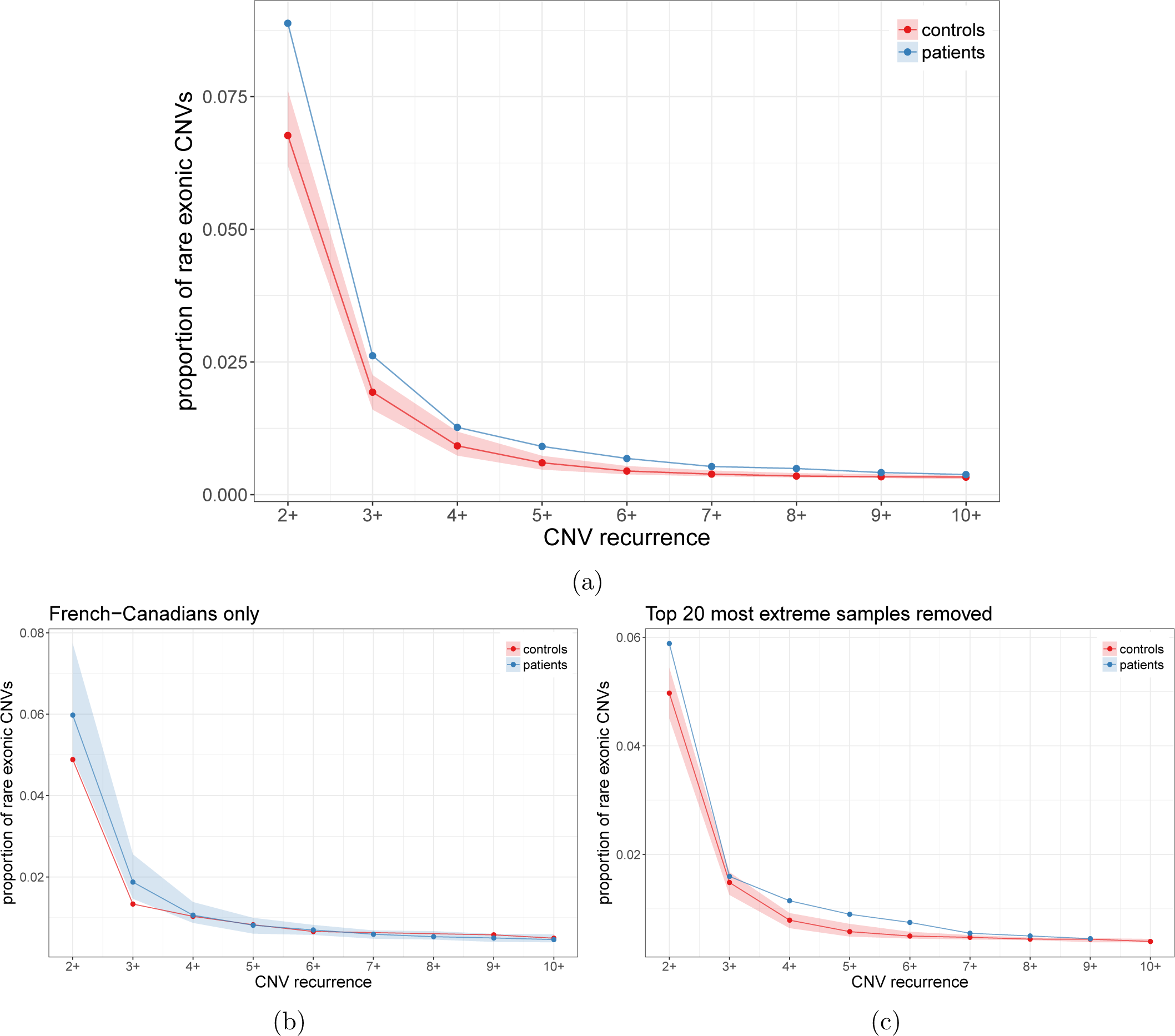
Rare exonic CNVs are less private in the epilepsy cohort. Proportion of rare exonic CNVs (y-axis) seen in X or more individuals (x-axis). The ribbon shows the 5%-95% confidence interval. In b), only French-Canadians individuals were analyzed and we down-sampled the epilepsy cohort to match the sample size of the French-Canadians controls. In c), the top 20 samples with the most non-private rare exonic SVs were removed.

**Fig S13:**
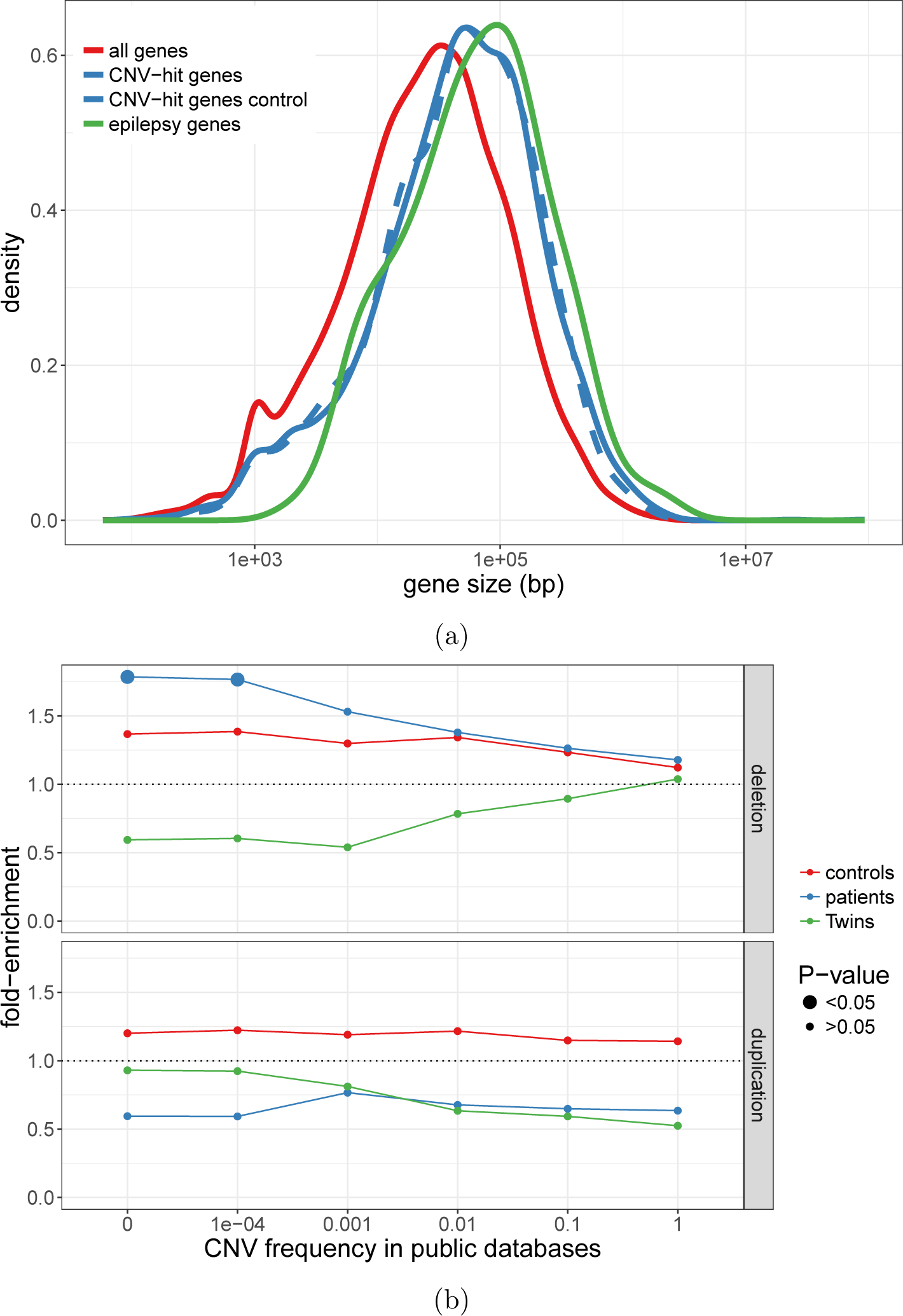
Enrichment in epilepsy genes. a) Epilepsy genes (red) are genes known to be associated with epilepsy. The control genes (dotted blue) are random genes selected so that the size distribution is similar to the sizes of genes hit by CNVs (plain blue). b) In three different datasets (color), genes hit by rare deletion (top) or duplications (bottom) at different frequency thresholds (x-axis) were tested for enrichment in epilepsy genes (y-axis, point-size).

**Fig S14:**
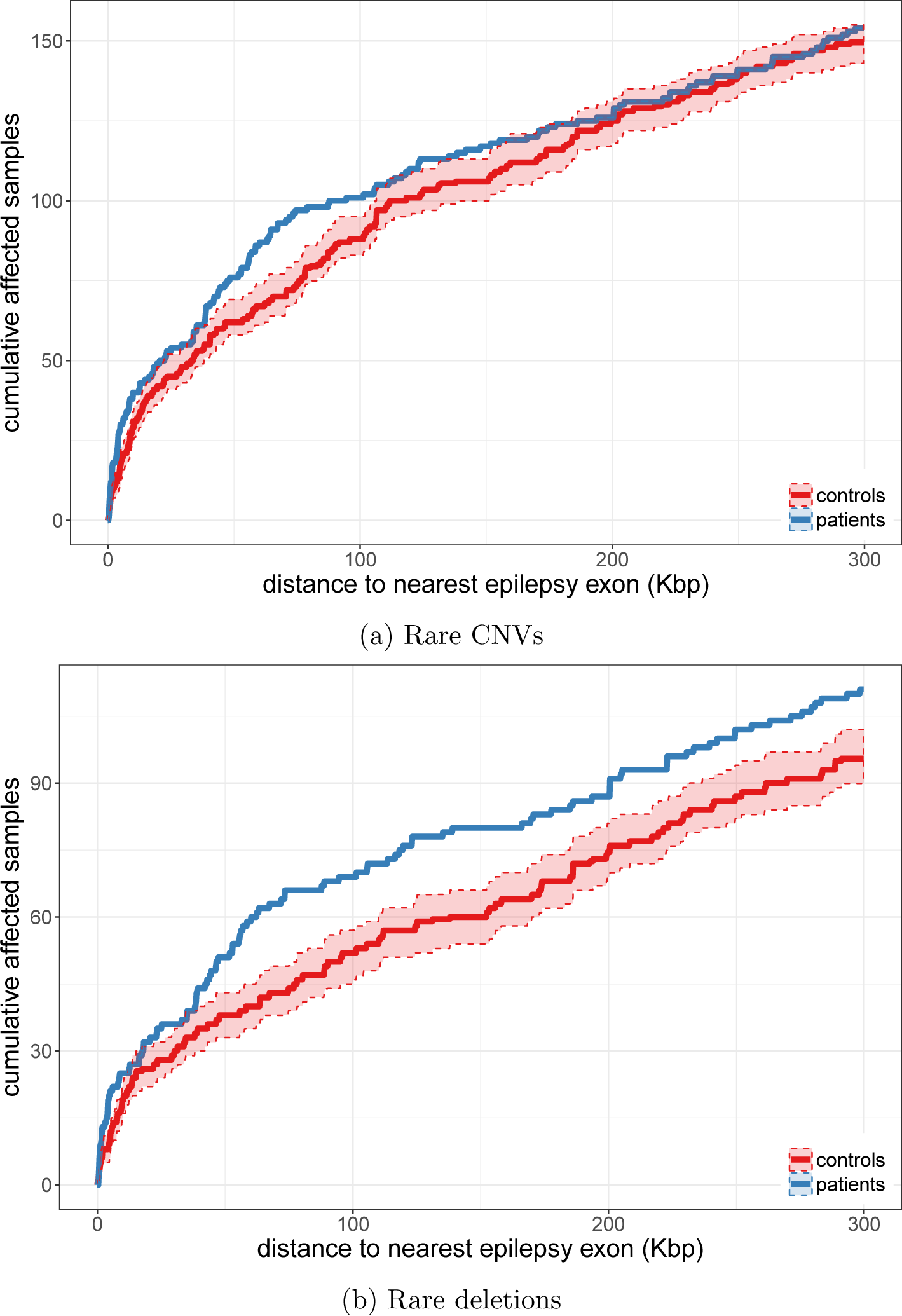
Rare non-coding CNVs near epilepsy genes. The graphs show the cumulative number of individuals (y-axis) with a rare non-coding variants located at X Kbp or less (x-axis) from the exonic sequence of a known epilepsy gene. The controls were down-sampled to the sample size of the epilepsy cohort. The ribbon shows the 5%/95% confidence interval. In a), deletions and duplications were considered; in b), only deletions were used.

**Fig S15:**
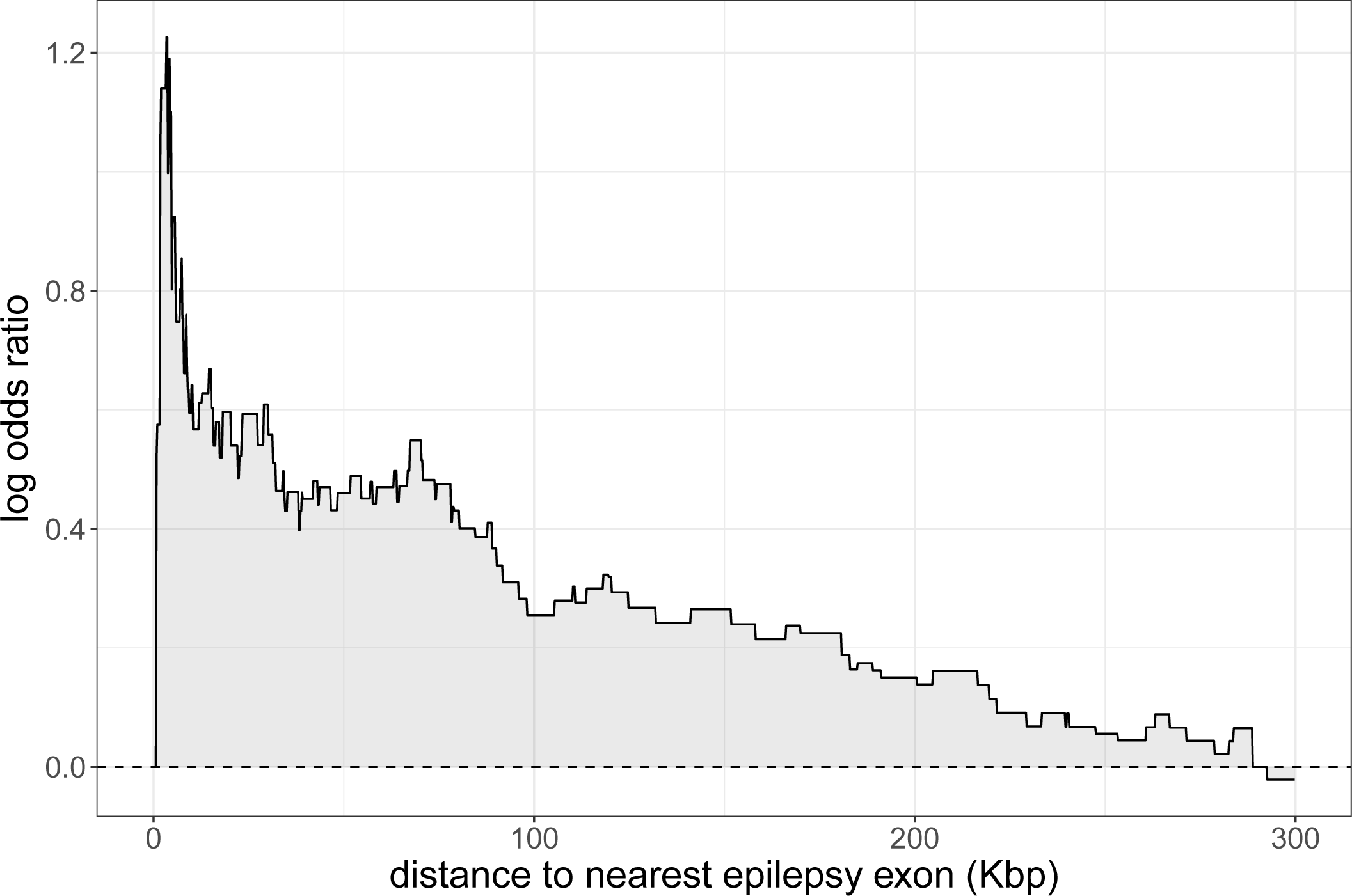
The enrichment in rare non-coding CNVs overlapping functional regions increases close to epilepsy genes. The graph shows the log odds ratio of having a rare non-coding CNV located at X Kbp or less (x-axis) from the exonic sequence of a known epilepsy gene. The y-axis shows the log odds ratio between epilepsy patients and controls. The controls were down-sampled to the sample size of the epilepsy cohort. We used CNVs overlapping regions functionally associated with the epilepsy gene (eQTL or promoter-associated DNase site).

**Fig S16:**
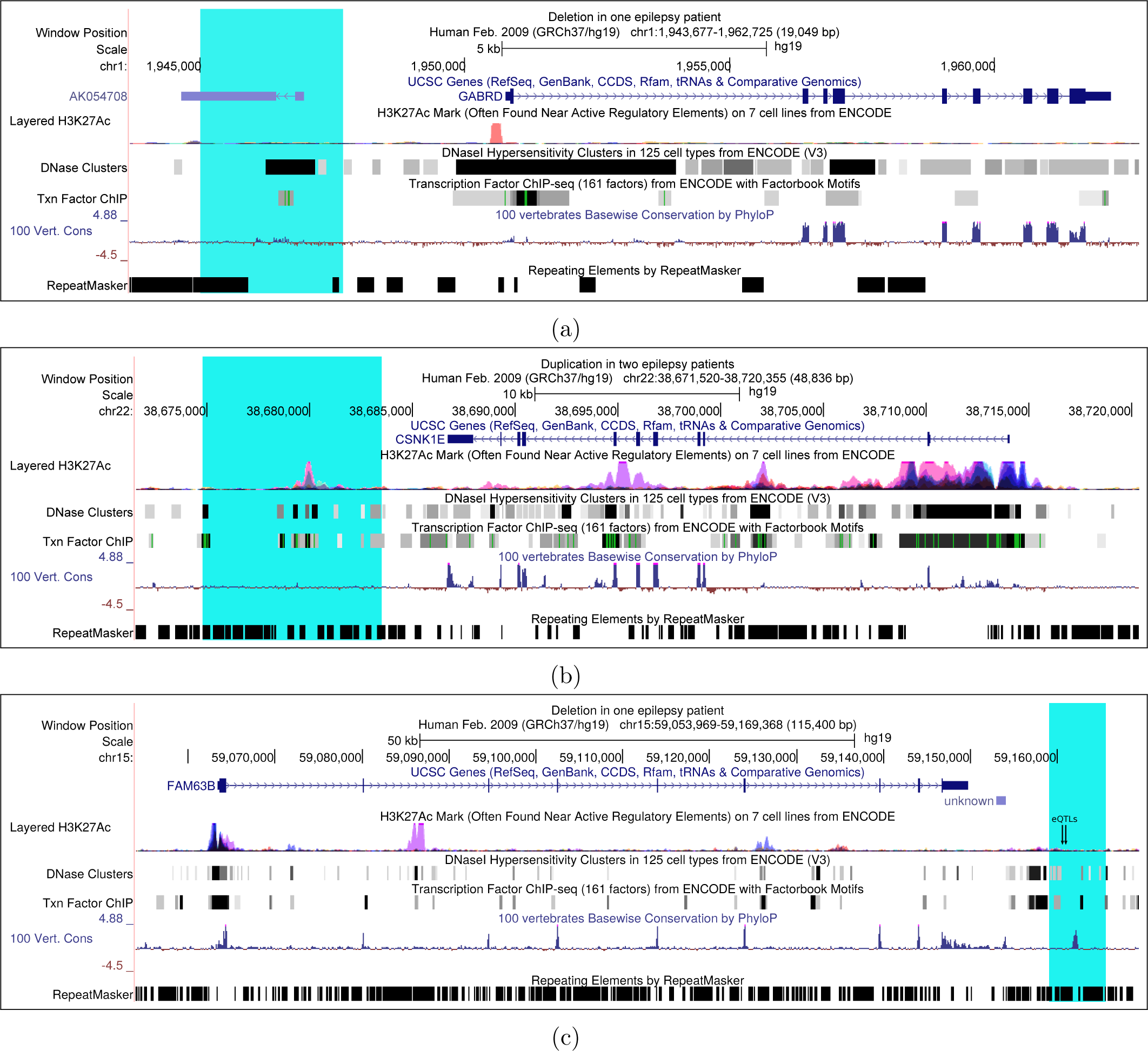
Non-coding CNVs with putative pathogenicity. a) 2.7 Kbp deletion in an epilepsy patient, never seen in controls or CNV databases. Three other epilepsy patients have a rare non-coding deletions located at less than 200 Kbp from the *GABRD* gene. b) 8.8 Kbp duplication in two epilepsy patients, never seen in controls or CNV databases and overlapping a regulatory region associated with *CSNK1E*. c) 6.5 Kbp deletion of an ultra-conserved regions downstream of *FAM63B*. Two expression QTLs for this gene are highlighted with arrows.

**Fig S17:**
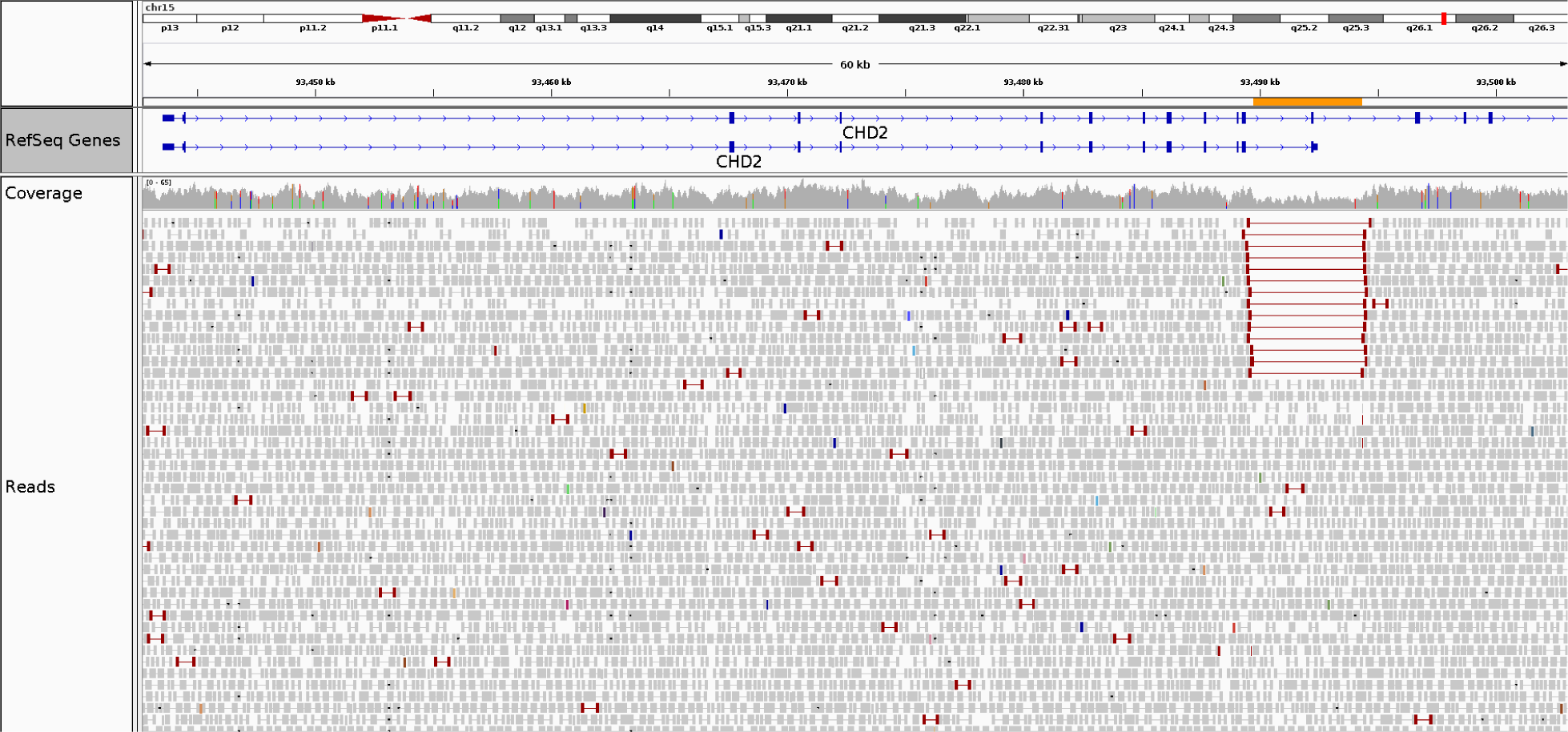
Small deletion of exon 13 in *CHD2*. Abnormal mapping of the read pairs highlighted in red support the deletion detected by PopSV using the read coverage. The deletion region is highlighted in orange.

**Fig S18:**
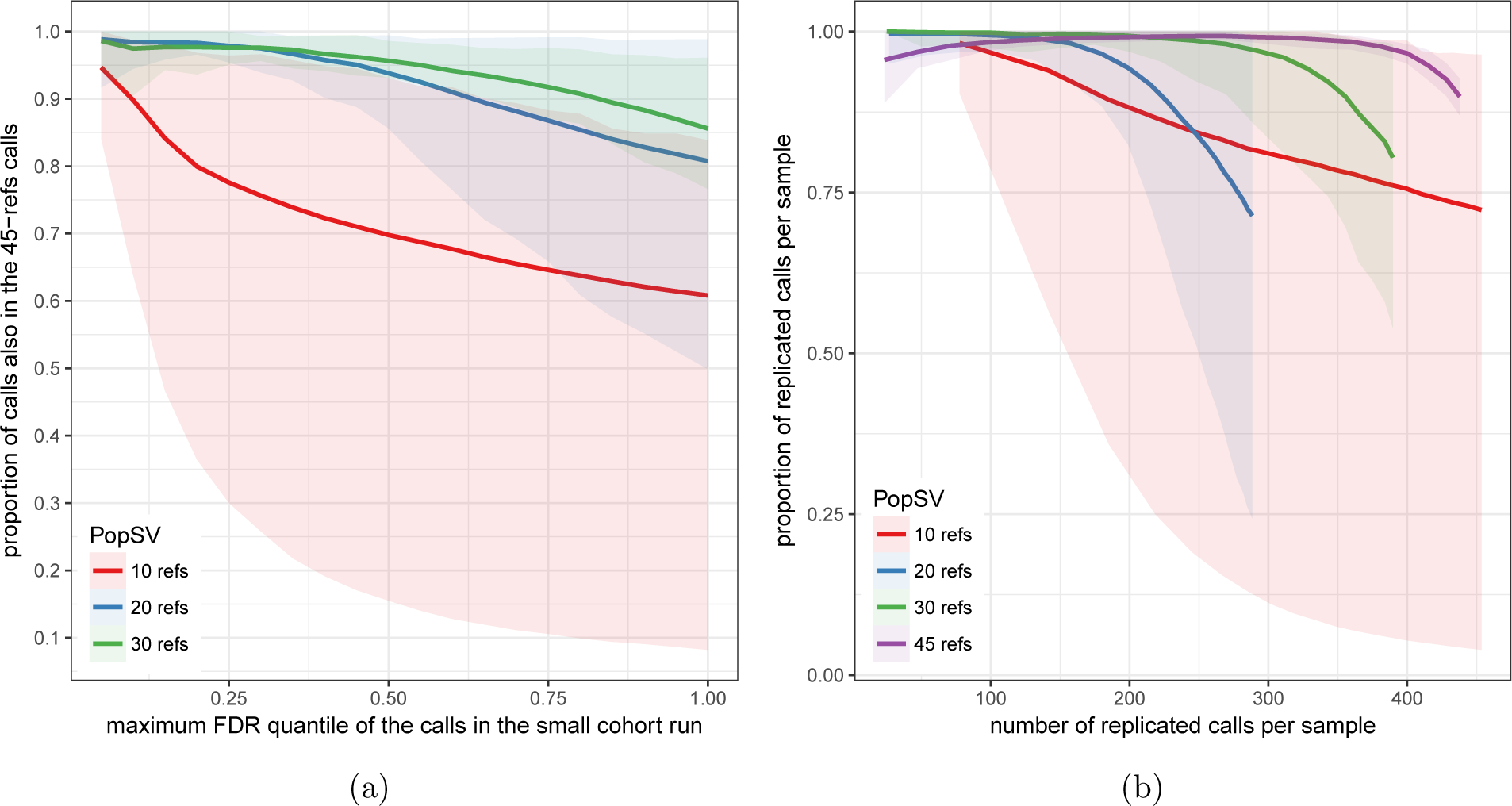
Reference cohort size and CNV detection quality. PopSV was run on the Twins study using 10, 20, 30 or 45 samples as reference (color). In a), the y-axis shows how many calls from the down-sampled run were found in the original 45-refs run. The x-axis represents the FDR threshold (lower threshold being more stringent). b) Replication in twins. For different cohort sizes and FDR thresholds, the number (x-axis) and proportion (y-axis) of calls replicated in the other twin is shown. In both graphs, the lines represents the median per sample and the ribbon the minimum/maximum value

**Fig S19:**
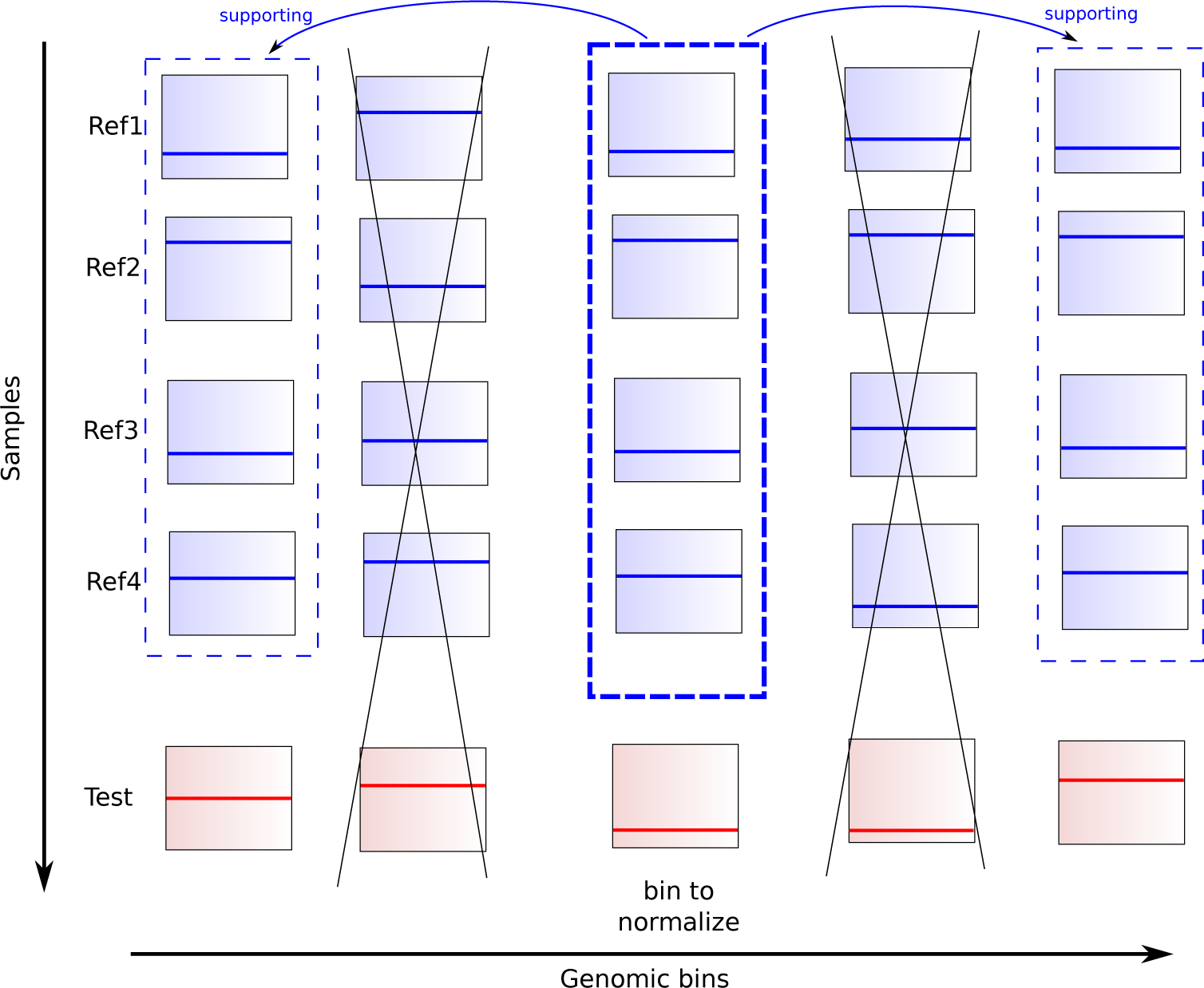
Targeted normalization. The coverage across the reference samples (blue) in the bin to normalize is used to find supporting bins across the genome. These supporting bins only are used to compute the normalization factor. The same supporting bins will be used to normalize the bin count in a test sample (red).

## 8 Supplementary Information

### 8.1 Epilepsy patients and sequencing

#### Ethics and patients recruitment

CENet is a Genome Canada and Genome Québec funded initiative that aims to bring personalized medicine in the treatment of epilepsy. Patients were recruited through two main recruitment sites at the Centre Hospitalier Universitaire de Montréal (CHUM) and the Sick Kids Hospital in Toronto. This study was approved by the Research Ethics Board at the Sick Kids Hospital (REB number 1000033784) and the ethics committee at the Centre Hospitalier Universitaire de Montréal (project number 2003-1394,ND02.058-BSP(CA)). Before their inclusion in this study, patients had to give written informed consents. The main cohort of this study was constituted of 198 unrelated patients with various types of epilepsy; 85 males and 113 females. The mean age at onset of the disease for our cohort was 9.2 (*±*6.7). Supplementary Table S1 presents a detailed description of the clinical features for the various individuals recruited in this study. DNA was extracted from blood DNA exclusively. 301 unrelated healthy parents of other probands from CENet were also included in this study and used as controls.

#### Libraries preparation and sequencing

gDNA was cleaned up using ZR-96 DNA Clean & Concentrator^TM^-5 Kit (Zymo) prior to being quantified using the Quant-iT^TM^ PicoGreen® dsDNA Assay Kit (Life Technologies) and its integrity assessed on agarose gels. Libraries were generated using the TruSeq DNA PCR-Free Library Preparation Kit (Illumina) according to the manufacturer’s recommendations. Libraries were quantified using the Quant-iT^TM^ PicoGreen® dsDNA Assay Kit (Life Technologies) and the Kapa Illumina GA with Revised Primers-SYBR Fast Universal kit (Kapa Biosystems). Average size fragment was determined using a LabChip GX (PerkinElmer) instrument.

The libraries were first denatured in 0.05N NaOH and then were diluted to 8pM using HT1 buffer. The clustering was done on a Illumina cBot and the flowcell was run on a HiSeq 2500 for 2×125 cycles (paired-end mode) using v4 chemistry and following the manufacturer’s instructions. A phiX library was used as a control and mixed with libraries at 1% level. The Illumina control software was HCS 2.2.58, the real-time analysis program was RTA v. 1.18.64. Program bcl2fastq v1.8.4 was then used to demultiplex samples and generate fastq reads. The average coverage was 37.6x *±* 5.6x. The filtered reads were aligned to reference Homo sapiens assembly b37. Each readset was aligned using BWA [59] which creates a Binary Alignment Map file (.bam). Then, all readset BAM files from the same sample are merged into a single global BAM file using Picard. Insertion and deletion realignment was performed on regions where multiple base mismatches were preferred over INDELs by the aligner since it appears to be less costly for the algorithm. Such regions were found to introduce false positive variant calls which may be filtered out by realigning those regions properly. Once local regions were realigned, the read mate coordinates of the aligned reads needed to be recalculated since the reads are realigned at positions that differ from their original alignment. Fixing the read mate positions is performed using Picard. Aligned reads were marked as duplicates if they have the same 5’ alignment positions (for both mates in the case of paired-end reads). All but the best pair (based on alignment score) were marked as a duplicate in the `.bam file`. Duplicates reads were excluded in the subsequent analysis. Marking duplicates was performed using Picard.

### 8.2 Testing for technical bias in WGS

To investigate the bias in read depth (RD), we first fragmented the genome in non-overlapping bins of 5 Kbp. The number of properly mapped reads was used as RD measure, defined as read pairs with correct orientation and insert size, and a mapping quality of 30 (Phred score) or more. In each sample, GC bias was corrected by fitting a Loess model between the bin’s RD and the bin’s GC content. Using this model, the correction factor for each bin was estimated from its GC content. Bins with extreme coverage were identified when deviating from the median coverage by more than 3 standard deviation. After these conventional intra-sample corrections, RD across the different samples were combined and quantile normalized. At that point the different samples had the same global RD distribution and no bins with extreme coverage or GC bias. Two control RD datasets were constructed to represent our expectation when no bias is present. One was derived from the original RD by shuffling the bins’ RD in each sample. In the second, RD was simulated from a Normal distribution with mean and variance fitted to the real distribution. Simulation or shuffling ensures that no region-specific or sample-specific bias remains. To investigate region-specific bias, we computed the mean and standard deviation of the RD in each bin across the different samples. The same was performed in the control datasets. If there is no bias, the distribution of these estimators should be similar in the original, shuffled and simulated RD. Next, to investigate experiment-specific bias, we retrieved which sample had the highest coverage in each bin. Then we computed, for each sample, the proportion of the genome where it had the highest coverage. If no bias was present, e.g. in the shuffled and simulated datasets, each sample should have the highest coverage in 100/N % of the genome (with N the number of samples). If an experiment was more affected by technical bias, it would be more often extreme. The same analysis was performed monitoring the lowest coverage.

The same analysis was ran after correcting the coverage in the Twin dataset using the QDNAseq pipeline [40]. The reads were counted in 5 Kbp bins using the function `binReadCounts`. GC bias and mappability were corrected using the following functions (with default parameters): `applyFilters`, `estimateCorrection`, `correctBins`, `normalizeBins`, `smoothOutlierBins`.

### 8.3 PopSV

#### Binning and coverage measure

The genome is fragmented in non-overlapping consecutive bins of fixed size (5 Kbp). In each bin and each sample the number of reads that overlap the bin and are properly mapped are counted to get a measure of coverage. Read pairs with correct orientation and insert size, and a mapping quality of 30 (Phred score) or more are considered properly mapped. The bin counts were then corrected for GC bias. In each sample, a LOESS model was fitted between the bin’s count and bin’s GC content. A normalization factor was then defined for each bin from its GC content.

#### Constructing the set of reference samples

In the epilepsy study and the Twins dataset we used all the samples as reference. In the renal cancer dataset we used the normal samples as reference. For each dataset, a Principal Component Analysis (PCA) was performed across samples on the counts normalized globally (median/variance adjusted). The resulting first two principal components are used to verify the homogeneity of the reference samples. In the presence of extreme outliers or clear sub-groups, a more cautious analysis would be recommended. For example, outliers can remain in the set of reference samples but flagged as they might potentially harbor more false calls later. Independent analysis in each of the identified sub-group is also a solution, especially when the same samples are to be used as reference. Although our three datasets showed different levels of homogeneity, we did not need to exclude samples or split the analysis. The effect of weak outlier samples was either corrected by the normalization step or integrated in the population-view. Moreover, the principal components were used to select one control sample from the final set of reference samples. This sample is used in the normalization step as a baseline to normalize other samples against. We picked the sample closest to the centroid of the reference samples in the Principal Component space.

#### Normalization

Although uniformity of the coverage across the genome is not required for our approach, RD values must be comparable across samples. When a particular region of the genome is tested, sample specific variation of technical origin must be minimized. This is done through a normalization step. Naive global normalization approaches like the Trimmed-Mean M(TMM) or quantile normalization have been first implemented and tested. The TMM normalization robustly aligns the mean RD value in the samples. Quantile normalization forces the RD distribution to be exactly the same in each sample. After witnessing the presence of uncharacterized sample-specific variation, we implemented a more suited normalization. Targeted normalization uses information across the set of reference samples to identify similar bins across the genome and normalize their counts separately (Figure S19). For each bin, the top 1000 bins with similar coverage patterns across the reference samples are used to normalize the coverage of the bin. TMM normalization is used on these top 1000 bins to derive the correct normalization factor for the bin to normalize. Similarity between two bins is measured using Pearson correlation between the counts across the reference samples. Hence, the top 1000 bins are most similar in term of relative coverage across the samples to the coverage in the bin to normalize. If some bias is present in some samples, the top 1000 bins should also harbor this bias. Hence, other regions with similar bias patterns are used to correct for it. In this targeted approach, each genomic region is normalized independently. The 1000 supporting bins are saved and used to normalized new samples (e.g. case sample). Although computationally expensive, it ensures that all bins are normalized with the same effort. In contrast, global normalization or even PCA-based approaches corrects for the most common or spread bias, but a subset of regions with specific bias might not be corrected. In order to compare the performance of the different normalization approaches we computed a set of quality metrics. The normalized RD will need to be suited for testing abnormal pattern across samples: under the null hypothesis, i.e. for normal bins, the RD should be relatively normally distributed and the samples rank should vary randomly from one bin to the other. The first metric is the proportion of bins with non-normal RD across the samples. Shapiro test was performed on each bin and a P-value lower than 0.01 defined non-normal RD. Then, the randomness of the sample ranks was tested by comparing the RD of each sample a region with the median across all samples. In regions of 100 consecutive bins, we counted how many times the RD in a sample was higher than the median across sample. If the ranks are random, this value should be around 0.5. The probability under the Binomial distribution is computed for each sample and corrected for multiple testing using Bonferroni correction. If any sample has an adjusted P-value lower than 0.05, we consider that the region has non-random ranks. The resulting QC metric is simply the proportion of regions with non-random sample ranks. This QC is specifically testing how much sample-specific bias remains. The remaining QC metrics look at the Z-score distribution in each sample. The proportion of non-normal Z-scores is computed by comparing the density curves of the Z-scores and simulated Normal Z-scores. We compute the proportion of the area under the density curve that does not overlap the Normal density curve. This estimate of the proportion of non-normal Z-scores is computed in each sample. The final metrics are the average and maximum across the samples.

#### Abnormal RD test and Z-score computation

The test is based on Z-scores computed for each bin, corrected afterward for multiple testing. The Z-score represents how different the read count in the tested sample is from the reference samples. It is simply: 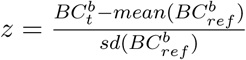 where 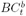 is the bin count, i.e. the number of reads, in bin *b* and sample *t*. Inevitably some samples are hosting common CNVs. We observed that just a couple of samples hosting a CNVs could be enough to bias the standard deviation used in the score computation and mask these CNVs in the coming tests. In many cases the RD signal was clearly showing several groups of samples with proportional read counts. To improve the Z-score computation in those regions, a simple approach was used: the samples were stringently clustered using their RD and the group with higher number of samples was chosen as reference and used to compute the mean and standard deviation for the Z-score computation. In practice, this clustering affects only bins with clear clusters but would remove just a few or no samples in most situations. Furthermore, a median-based estimator was used for the standard deviation as it is less sensitive to outlier removal. A trimmed mean was also preferred over normal mean for its robustness to outliers.

#### Significance and multiple testing correction

The Z-scores for all the bins of a sample are pooled and significance is estimated. Under the null hypothesis of normally distributed read counts, the Z-scores should also follow a normal distribution. For multiple testing correction, the Z-score empirical distribution is used to fit a normal and estimate the P-value and Q-value of each test. This step is performed using fdrtool R package. By default, the null distribution fitting for P-value computation assumes that only a low proportion of bins violates the null hypothesis. In aberrant genomes, e.g. in tumor samples, it is often an unrealistic assumption. We devised a new strategy to set the proportion of the empirical distribution, later used to estimate the null distribution variance. Here the null Z-score distribution is assumed to be centered on 0 and its variance is estimated by trimming the tails of the empirical distribution. To find a correct trimming factor, an iterative approach started from a low trimming factor and increased its value until reaching a plateau for the variance estimator. Indeed, once the plateau is reached, additional trimming does not change the estimated variance because there is no more abnormal Z-scores, only the central part of the null distribution. Samples with an important proportion of abnormal genome, e.g. tumor samples, showed more appropriate fit. Of note, the P-values for positive Z-scores (duplication) and negative Z-scores (deletion) are estimated separately. Thus, imbalance in the deletion to duplication ratio, or large aberration that lead to asymmetrical Z-score distribution does not affect the P-value estimation. Multiple testing correction is performed after pooling all the P-values.

#### Segmentation, copy number estimation and other metrics

Following the significance estimation, consecutive bins with abnormal coverage are merged into a call. Consecutive or nearby abnormal bins (e.g. one bin size apart) are merged into one variant if in the same direction (deletion or duplication). In PopSV’s R package, the P-values can also be segmented using circular binary segmentation [60].

In addition to the Z-score, P-value, Q-value and number of bins of each call, PopSV retrieves the average coverage in the reference samples and the fold change in the sample tested. The copy number is estimated by dividing the coverage in a region by the average coverage across the reference samples, multiplied by 2 (as diploidy is expected). In our bin setting, the estimation is correct if the bin spans completely the variant. For this reason we trust the copy number estimate for calls spanning 3 or more consecutive bins, as it is computed using the middle bin(s) which completely span the variant. In other cases we expect the copy number estimate to be under-estimated. All this additional information can be used to order or retrieve high confidence calls. For examples, several consecutive bins or a copy number estimate around an integer value increases our confidence in a call. In our benchmark, we used the entire set of calls.

### 8.4 Validation and benchmark of PopSV

We compared PopSV to FREEC [10], CNVnator [9] and cn.MOPS [11], three popular RD methods that can be applied to WGS datasets to identify CNVs. FREEC segments the RD values of a sample using a LASSO-based algorithm while CNVnator uses a mean-shift technique inspired from image processing. cn.MOPS considers simultaneously several samples and detects copy number variation using a Poisson model and a Bayesian approach. We also ran LUMPY [8] which uses an orthogonal mapping signal: the insert size, orientation and split mapping of paired reads.

FREEC and CNVnator were run on each sample separately, starting from the BAM file. FREEC internally corrects the RD for GC and mappability bias. In order to compare its performance across the entire genome, the minimum telocentromeric distance was set to 0. The remaining parameters were set to default. Of note an additional run with slightly looser parameter (‘breakPointThreshold=0.6’) was performed to get a larger set of calls used in some parts of the in silico validation analysis to deal with borderline significant calls. CNVnator also corrects internally for GC bias. We used default parameters. For the analysis using higher confidence calls, we used calls with either ‘eval1’ or ‘eval2’ lower than 10-5 (instead of the default 0.05). cn.MOPS was run on the same GC-corrected bin counts used for PopSV. All the samples are analyzed jointly. Of note an additional run with slightly looser parameter (’upperThreshold=0.32’ and ‘lowerThreshold=-0.42’) was performed to get a larger set of calls used in some parts of the in silico validation analysis to deal with borderline significant calls. For LUMPY, the discordant reads were extracted from the BAMs using the recommended commands. Split-reads were obtained by running YAHA [62] with default parameters. All the CNVs (deletions and duplications) larger than 300 bp were kept for the upcoming analysis. Calls with 5 or more supporting reads were considered high-confidence.

First, we compared the frequency at which a region is affected by a CNV using the calls from the different methods. In order to investigate how many systematic calls are present in a typical run, we compare the frequency distributions on average per sample. In figure S4, the bars represents the average proportion of a sample’s calls in each frequency range.

Then, the samples were clustered using the CNV calls. The distance between two samples A and B is defined as : 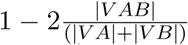 where VA represents the variants found in sample A, VAB the variants found in both A and B, and —V— the cumulative size of the variants. Hence, the similarity between two samples is represented by the amount of sequence called in both divided by the average amount of sequence called. This distance is used for hierarchical clustering of the samples. Different linkage criteria (“average”, “complete” and “Ward”) were used for the exploration. In our dendograms we used the “average” linkage criterion. The same clustering was performed using only calls in regions with extremely low coverage (reference average i 10 reads).

To assess the performance of each method, we measured the number of CNVs identified in each twin that were also found in the matching twin. In order to avoid missing calls with borderline significance, we used slightly less confident calls for the second twin. We removed calls present in more than 50% of the samples to ensure that systematic errors were not biasing our replication estimates. Hence, a replicated call is most likely true as it is present in a minority of samples but consistently in the twin pair. Even if we removed systematic calls, the most frequent calls in the cohort are more likely to look replicated by chance, compared to rare calls. To normalize for this effect, we use the frequency distribution to compute the number of replicated calls expected by chance. In practice the null concordance for each call is simulated by a Bernoulli distribution of parameter the frequency of the call. This number of replicated calls by chance is subtracted to the original number of replicated calls to give an adjusted measure of sensitivity. Although we do not know the true number of variant, this number of replicated calls is used to compare the different methods. When possible, the low-quality calls were also gradually filtered to explore the effect on the replication metrics. For CNVnator, we used the minimum of the eval1 and eval2 columns, with lower values corresponding to higher quality calls. For LUMPY, the number of supporting reads was used. For PopSV, we filtered calls based on adjusted P-values.

In addition to their replication, we compared which regions were called by several methods. For each of the calls found in less than 50% of the samples, we overlapped the region with calls from other methods in the same sample. If calls from another method overlapped we considered the call shared and saved which methods shared the call. To focus on on high quality calls we considered calls found by at least two methods and computed the proportion of calls from one method found by each of the other methods. This metric captures how much each method recovers high-quality calls from a second method.

#### Concordance between different bin sizes

We compared calls using small bins (500 bp) and calls using larger bins (5 Kbp). In theory, calls from the 5 Kbp analysis should be supported by many 500 bp calls. We also expect large stretches of 500 bp calls to be detected in the 5 Kbp analysis. This comparison is informative as it explores the quality of the calls, the size of detectable events and the resolution for different bin sizes. First we counted how many small bin calls supported any large bin call. These metrics were separated according to the size of the large bin call. Overall, we find that 5 Kbp calls are well supported by 500 bp calls, with only 14% of the 5 Kbp bins not supported by any 500 bp bin (Figure S9a). To investigate large bin calls with no supporting small bin call, we display the average Z-scores in the small bins overlapping large bin calls to test if the lack of support is due to lower confidence or real discordancy between the different runs. If the Z-scores in the small bins deviates from 0 in the correct direction, we conclude that they support the large bin call. Even for these unsupported 5 Kbp calls, we find that the 500 bp bins RD was consistently enriched (or depleted) although not enough to be called with confidence (Figure S9b and S9c). This is expected given the higher background noise in the 500 bp analysis that will reduce the power to call these variants. Next, we looked at the proportion of 500 bp calls, grouped by size, that were found in the 5 Kbp calls. More specifically, we grouped them by size to verify that large enough small bin calls are present in the large bin calls. This analysis is used to both test the sensitivity of PopSV with a particular bin size, and its resolution to variants smaller than the bin size. Indeed, this framework allow us to ask questions such as: how much of the variants spanning only half a bin are detected ? We find that the concordance gradually increases until the 500 bp calls reach 5 Kbp in size where the concordance rises to nearly 100% (Figure S9d). This suggests that PopSV is able to detect approximately 75% of the events as large as half its bin size, and almost all events larger than its bin size. As expected, only a small proportion of the small 500 bp calls overlap 5 Kbp calls and they likely corresponds to fragmented larger calls. Considering the trade-off between bin size and noise, this suggests running PopSV with a few bin sizes to better capture variants of different sizes.

### 8.5 CNV detection in the CENet cohorts

CNVs were called using PopSV using 5 Kbp bins and all the samples from both the epilepsy and control cohorts as reference. We annotated the frequency of the CNVs using germline CNV calls from the Twin and cancer datasets (internal database) as well as four public CNV databases:

- CNVs from Phase 1 of the 1000 Genomes Project as identified by Genome STRiP [13].
- SV from the 1000 Genomes Project phase 3 [45].
- Genome of Netherlands [44].
- CNVs from the Simons Genome Diversity Project [46].

CNVs were annotated with the maximum frequency in the databases. For each CNV to annotate, any overlapping CNV in the CNV databases were considered. This is a stringent criterion that ensures that the entire regions of a rare CNV, for example, is never affected by common CNVs in the databases. Hence, a rare CNV is defined as present in less than 1% of the samples in each of the five CNV databases.

### 8.6 CNV enrichment in exonic region and around epilepsy genes

#### Enrichment in exons

For each cohort, we retrieved the CNV catalog by merging CNV that are recurrent in multiple samples. Hence, the CNV catalog represents all the different CNVs found in each cohort. To control for the population size, we sub-sampled 150 samples in each cohort a hundred times. For each sub-sampling and each cohort, control regions are selected to fit the size distribution of the CNV catalog and the overlap with centromere, telomeres and assembly gaps (details in the next section).

Then, we computed the proportion of CNV and control regions that overlap an exon. The fold-enrichment is the ratio of these proportions and represents how much more/less of the CNVs overlap an exon compared to the control regions. The boxplot in Figure 2c shows the distribution of the 100 sub-sampling in each cohort.

To test if the difference observed between the cohort is significant, the *cohort* labels were permuted 10,000 times and the difference in median across the 100 sub-sampling was saved. The resulting P-value was computed as 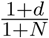 where *d* is the number of times the permuted difference was greater or equal to the observed difference, and *N* is the number of permutations.

The same analysis was repeated for exons from genes with a probability of loss-of-function intolerance [47] higher than 0.9. These genes were called *LoF intolerant genes* in Figure 2c. Small (i 50Kbp) and large (¿50 Kbp) CNVs were analyzed separately. The analysis was repeated using rare CNVs only.

#### Selecting control regions

The control regions must have the same size distribution as the regions they are derived from (e.g. CNVs in a CNV catalog). We also controlled for the overlap with centromere, telomeres and assembly gaps (CTGs) to avoid selecting control regions in assembly gaps where no CNV or annotation is available. To select control regions, thousands of bases were first randomly chosen in the genome. The distance between each base and the nearest CTG was then computed. At this point, selecting a region of a specific size and with specific overlap profile can be done by randomly choosing as center one of the bases that fit the profile:

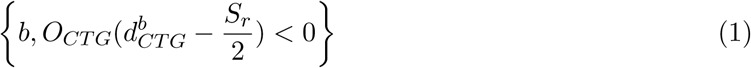

with *O*_*CTG*_ equals 1 if the original region overlaps with a CTG, −1 if not; 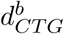 is the distance between base *b* and the nearest CTG; and *S*_*r*_ is the size of the original region. For each input region, a control region was selected and had by construction the exact same size and overlap profile.

#### Recurrence of rare exonic CNVs

In each cohort, we retrieved the CNV catalog of rare (¡1% in all 5 public datasets) exonic CNVs. We annotated each CNV with its recurrence in the cohort. We then evaluated the proportion of the CNVs in the catalog that are private (i.e. seen in only one sample), or seen in X samples or more. This cumulative proportion of CNVs is shown in Figure S12a. The control cohort was down-sampled a thousand times to the same sample size as the epilepsy cohort. These down-sampling provided a confidence interval (ribbon in Figure S12a) and an empirical P-value.

We performed the same analysis after removing the top 20 samples with the most non-private rare exonic CNVs (Figure S12b). With this analysis, we tried to remove the potential effect of a few extreme samples.

We also repeated the analysis using only French-Canadians individuals, to ensure that the observed differences are not caused by rare population-specific variants (Figure S12b).

### CNVs and epilepsy genes

We used the list of genes associated with epilepsy from the EpilepsyGene resource [48] which consists of 154 genes strongly associated with epilepsy. For a particular set of CNV we count how many of the genes hit are known epilepsy genes. We noticed that the epilepsy genes tend to be large, and genes hit by CNVs also (Figure S13a). This could lead to a spurious association so we also performed a permutation approach that controls for the size of the genes. To control for the gene size of epilepsy genes and CNV-hit genes, we randomly selected genes with sizes similar to the genes hit by CNVs and evaluated how many of these were epilepsy genes. After ten thousand samplings, we computed an empirical P-value (see Supplementary Information). The permutation P-value was computed as 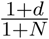 where *d* is the number of times the number of epilepsy genes in the random set of genes was greater or equal to the one in genes hit by CNVs, and *N* is the number of permutations. Using this sampling approach we tested different sets of CNVs: deletion or duplications of different frequencies in the epilepsy cohort, control individuals and samples from the twin study.

To investigate rare non-coding CNV close to known epilepsy genes, we counted how many patients have such a CNV at different distance thresholds. For example, how many patients had a rare non-coding CNV at 10 Kbp of an epilepsy gene’s exon or closer. We compared this cumulative distribution to the control cohort, after down-sampling it to the sample size of the epilepsy cohort. Down-sampling was also used to produce a confidence interval, represented by the ribbon in Figure 3c). This analysis was repeated using deletions only. Each epilepsy gene was also tested for an excess of rare non-coding deletions in patients versus controls using a Fisher test.

In order to retrieve non-coding CNV that might have a functional impact, we downloaded eQTLs associated with the epilepsy genes, as well as DNase 1 hypersensitive sites associated with the promoter of epilepsy genes. The eQTLs are provided by the GTEx project [50]. Pairs of associated DNase 1 hypersensitive sites and associated genes [51] were downloaded at http://www.uwencode.org/proj/Science_Maurano_Humbert_et_al/data/genomewideCorrs_above0.7_promoterPlusMinus500kb_withGeneNames_35celltypeCategories.bed8.gz.

A Kolmogorov-Smirnov test was used to compare the distance distributions in epilepsy patients versus controls. We also computed the odds ratio of having such a CNV for different distance thresholds between epilepsy patients and controls. For a distance *d*, we computed:

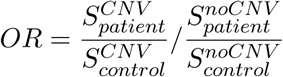

where 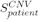 is the number of patients with a rare non-coding CNV overlapping a functional region and located at *d* bp or less from the exon of a known epilepsy gene.

